# WISH-BONE: Whole-mount In Situ Histology, to label osteocyte mRNA and protein in 3D adult mouse bones

**DOI:** 10.1101/2024.03.14.585053

**Authors:** Quentin A. Meslier, Timothy J. Duerr, Brian Nguyen, James R. Monaghan, Sandra J. Shefelbine

**Affiliations:** Department of Bioengineering, Northeastern University, Boston, MA, United States; Department of Biology, Northeastern University, Boston, MA, United States; LifeCanvas Technologies, Cambridge, MA, United States; Institute for Chemical Imaging of Living Systems, Northeastern University, Boston, MA, United States

## Abstract

Bone is a three dimensional and highly dynamic tissue under constant remodeling. Commonly used tools to investigate bone biology require sample digestion for biomolecule extraction or only provide two-dimensional spatial information. There is a need for 3D tools to investigate spatially preserved biomarker expression in osteocytes. In this work, we present a new method, WISH-BONE, to label osteocyte mRNA and protein in whole-mount mouse bone. For mRNA labeling, we used Hybridization Chain Reaction – Fluorescence *In Situ* Hybridization to label genes of interest in osteocytes. For protein labeling, samples were preserved using an epoxy-based solution that protects tissue structure and biomolecular components. Then an enzymatic matrix permeabilization step was performed to enable antibody penetration. Immunostaining was used to label various proteins involved in bone homeostasis. We also demonstrate the use of customized fluorescent nanobodies to target and label proteins in the cortical bone. However, the relatively dim signal observed from nanobodies staining limited detection. In this study, we share protocols, highlight opportunities, and identify the challenges of this novel 3D labeling method. They are the first protocols for whole-mount osteocyte 3D labeling of mRNA and protein in mature mouse bones. WISH-BONE will allow the investigation of molecular signaling in bone cells in their 3D environment and could be applied to various bone-related fields of research.

## Introduction

Bone is a mineralized tissue composed of an organic and an inorganic matrix that provides mechanical properties necessary for support, movement, and organ protection. Bone is also a highly dynamic tissue that is continuously remodeled, which is critical for fracture healing, mechanoadaptation, and calcium homeostasis. Alteration of bone structure or imbalance of bone remodeling can result in various pathologies such as osteoporosis. In addition, bone represents one of the most common sites of cancer metastasis. Research efforts in bone biology enable the identification of signaling pathways and potential targets for the development of therapeutics.

Quantitative polymerase chain reaction (qPCR) and RNA-sequencing are commonly used methods to quantify the bulk amount of mRNA expressed in bone samples. However, isolation of specific cell types from bone samples is challenging. Single-cell RNA sequencing enabled the identification of new bone-cell clusters within bone cell lineage and investigation of individual cell’s transcriptomes [1]. Quantifying mRNA requires extraction which results in loss of spatial context. In addition, osteocyte isolation is challenging due to their location deep in the bone matrix. Laser capture microdissection has been used to isolate specific regions of the bone and extract a more homogenous cell population [2], [3].

To gain 2D spatial information of gene expression, *in situ* hybridization and immunolabeling have been used on thin tissue sections to investigate mRNA and protein expression respectively in the bone. In addition, spatial transcriptomics is also a promising tool for bone research as it addresses spatial limitations of mRNA sequencing [4]. However, these different methods are performed on 2D bone sections. Careful selection of the region to be studied is necessary to observe the biological response between experimental conditions. In addition, the selection of a 5-10µm-thick region may not be representative of the overall biological response. Capturing the heterogeneity of the bone in 2D sections can be challenging for many applications. For example, callus formation and composition during bone fracture healing vary spatially [5]. Bone metastases disrupt normal bone geometry and increase heterogeneity of the tissue in specific locations [6]. Bone development varies along the different axes [7]. Some bone pathologies, such as fibrous dysplasia, locally affect bone structure and mechanical properties [8]. Abnormal fibrous tissues may be challenging to observe via traditional histology as it requires sectioning in the location of the abnormal bone. Finally, in the context of mechanoadaption, external forces drive bone adaptation. The osteogenic response depends on the spatially heterogeneous mechanical stimuli in the bone. Due to the inherent bone structure and 3D nature of the processes studied in bone, 2D methods are a limiting factor in bone research.

Whole-mount tissue clearing [9] enables imaging of tissue samples in 3D by removing light-absorbing and light-scattering elements, such as lipids, using organic or aqueous solutions. Some methods use additional crosslinker or encapsulation in hydrogels to protect samples’ biomolecules during clearing protocols [10]. For example, SHIELD (stabilization to harsh condition via intramolecular epoxide linkages to prevent degradation) [11] has been shown to preserve tissues structure during delipidation, while enabling protein labeling using antibodies.

Following clearing, a sample’s refractive index is matched to the mounting solution to reach optical transparency, which enables light penetration through the entire sample. Tissue clearing allows images to be acquired several millimeters deep; signal from noncleared tissues can only be acquired a few microns into the tissues using confocal or two-photon microscopy. More recently, tissue clearing protocols have been adapted for calcified tissue [12], [13], [14]. In bone, these methods have been used to image endogenous fluorescence from transgenic animals (no labeling needed) [12], [13], [15] and immunolabeling of elements in the bone canals, such as vascularization [16], [17], innervation [18], and the lymphatic system [19]. However, labeling of osteocyte mRNA and proteins in whole-mount bone remains challenging due to the osteocytes’ location within the dense bone matrix.

Osteocytes are the most abundant cell type in bone. They are embedded in the bone matrix and are known to play a critical role in the regulation of bone biology, including bone formation, resorption, and local mineral deposition. Osteocytes are also endocrine cells as they secrete factors that target other organs. Osteocytes are involved in fracture healing [20], [21] and high-incidence pathologies such as osteoporosis [22], but also rarer diseases such as hyperphosphatemia [23] and slcerosteosis [24]. Investigation of gene expression in osteocytes has provided valuable insight into bone biology by identifying new signaling pathways and new targets for therapeutic development [25], [26]. However, methods that allow investigation of the osteocytes gene expression in 3D remain to be developed.

In this work, we present WISH-BONE, a new method that enables labeling of mRNA and protein in 3D mouse bone. In addition to the detailed protocols, we share valuable insights on parameters that could affect results and potential applications for this method. WISH-BONE is opening new ways to investigate bone biology and will help to correlate biological responses to cells’ 3D environment.

## Methods

**Note**: In this study, all incubation was performed at room temperature and with shaking unless specified differently.

### Animal model

All animal experiments presented in this study were approved by Northeastern University’s Institutional Animal Care and Use Committee (IACUC). C57BL/6J mice of 18 to 23 weeks old were purchased from Jackson Laboratories. Mice were all used between the age of 20 weeks and 30 weeks old. They were housed in cages of 5, with 12-hour light and dark cycles, and under a regular diet.

### Sample collections, fixation, and decalcification

Mice were sacrificed via CO_2_ inhalation and cervical dislocation. Long bones and vertebrae were collected. External tissues were removed. The distal ends of the long bones were sectioned using a scalpel and bone marrow was flushed from the proximal end using a 1ml syringe filled with PBS. Samples were immediately placed in 5 ml of ice-chilled 4% paraformaldehyde (PFA) and kept for 24h at 4°C with shaking. The fixative solution was removed with 6×5min washes in 1X PBS. Bones were then placed in 5ml of 10% EDTA at 4°C with shaking for decalcification. EDTA solution was changed every 3 days for a total of 2 weeks. Once decalcified, samples were washed 6×5min in water.

**Note:** Samples for mRNA labeling were incubated with DEPC water and RNase-free solutions.

### Whole-mount Hybridization Chain Reaction -Fluorescence *In Situ* Hybridization (HCR-FISH)

Probes for whole-mount v3. HCR-FISH were generated as previously described [27], [28]. Briefly, probe designs were generated using the web-based tool Probegenerator (http://ec2-44-211-232-78.compute-1.amazonaws.com). 36 probe pairs for each gene were selected with preference for probes located first in the open reading frame, then the 5’ UTR, and finally the 3’ UTR. Probe designs are listed in supplementary table (Supplementals table 1). Oligonucleotide pools were ordered as lyophilized oPools (50 pmol) from Integrated DNA Technologies. Prior to use in whole-mount v3. HCR FISH, probes were resuspended in Tris-EDTA buffer at a concentration of 1 mM and stored at -20°C.

All mRNA labeling presented in this study was performed in the mouse tibia midshaft. The FISH protocol was adapted from Molecular Instruments and a previously described protocol [27]. After fixation and decalcification, midshafts were isolated using a scalpel. The midshaft region was defined by the proximal and distal tibia-fibula junctions. Fixed and decalcified samples were then dehydrated in iced cold methanol (MeOH) through an increasing graded series of incubations (25%, 50%, 75%, 100%, and 100% of MeOH) in PBS with 10% Tween (PBST), with 15 min incubations per step, and then stored overnight at -20°C. Samples were stored for up to 2 weeks before labeling. Samples were rehydrated using a decreasing concentration gradient of iced cold MeOH (75%, 50%, 25%) followed by incubation in 100% PBST. Tissues were permeabilized using proteinase K (10µg/ml) in PBS for 15 min. Post-fixation was performed using 4% PFA for 20 min and washed 3×5min in PBST. Samples were briefly washed using 30% hybridization buffer (Molecular Instruments, USA) for 5 min in at 37°C. Prehybridization was performed using hybridization buffer for 30 min at 37°C. Probes were diluted (1:200) in 30% hybridization buffer to a final concentration of 5 µM. Samples were placed in FISH probe solutions overnight at 37°C.

The following day, the probe solution was washed 4×15min using preheated 30% formamide wash solution (Molecular Instruments, USA) at 37°C, then washed for 5 min twice using 5x saline-sodium citrate with 10% Tween (5xSSCT). A pre-amplification step was performed by incubating samples in amplification buffer (Molecular Instruments, USA) for 30min. DNA hairpins (3µM stock solution) were heated at 90°C for 1 min 30 sec and added to amplification buffer with a dilution of 1:50 for a final concentration of 60 nM. Samples were incubated in hairpin solution overnight in the dark. After amplification, unbound hairpins were washed using 5xSSCT 4 times for 20 min. Nuclei were stained by incubating in Oxazole Yellow (1:1000) overnight at 4°C with shaking. Negative controls were incubated with DNA hairpins without prior incubation with probes.

### Whole-Mount Immunofluorescence, via antibodies

Following decalcification, samples were preserved using SHIELD (LifeCanvas Technologies, USA). SHIELD uses polyepoxy to crosslink amine groups at the biomolecules surface and protect their structure, preserving the physiological location of biomolecules and enhancing their mechanical stability [11]. Midshafts were isolated using a scalpel. The sectioning process was carried out at the two tibia-fibula junctions. SHIELD OFF Solution was prepared with 50% SHIELD-Epoxy, 25% SHIELD Buffer, and 25% water (LifeCanvas Technologies, USA). Tibia midshafts were incubated in 5ml of SHIELD OFF Solution overnight at 4°C with shaking. Next, samples were transferred to pre-heated SHIELD ON Buffer with 1% SHIELD Epoxy and incubated at 37°C overnight. Once samples were preserved, an enzymatic matrix permeabilization step was performed to enable antibody penetration, as illustrated in Figure 1. Collagenase type II produced by *Clostridium histolyticum* [29] was used to permeabilize the bone matrix.

**Figure 1:**
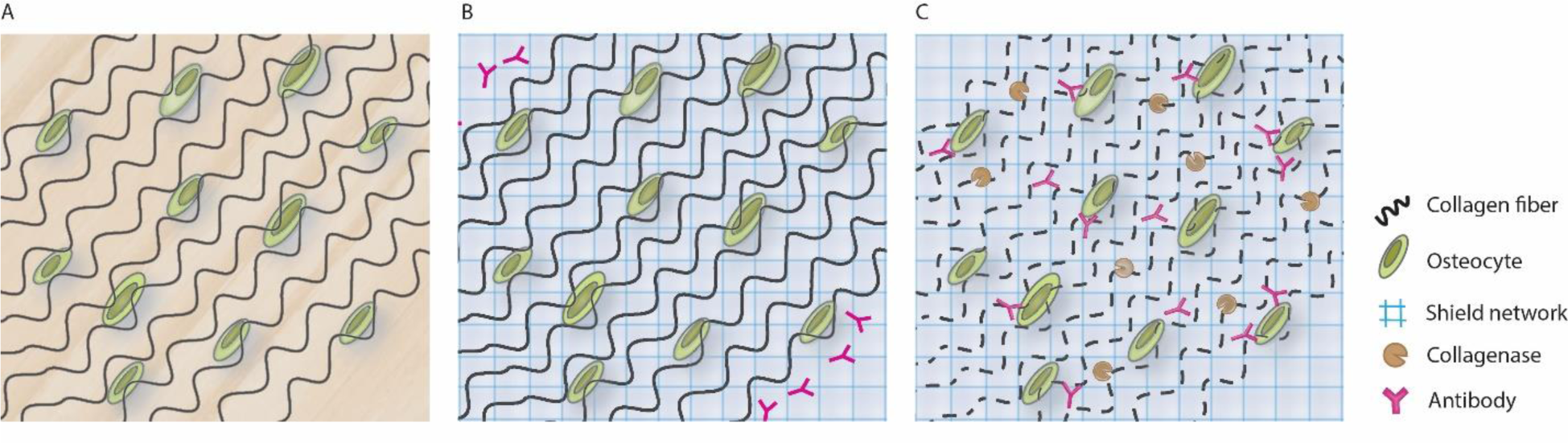
Method summary. A) Illustration of osteocytes embedded in the dense bone matrix. B) The sample is preserved using SHIELD (LifeCanvas Technologies, USA). C) Sample is permeabilized using collagenase allowing the penetration of the antibodies while limiting degradation of the samples.

Collagenase type II (C2-BIOC, Millipore Sigma) was rehydrated in 1x HBSS at a concentration of 10mg/ml and stored in 5 ml tubes at -20°C. Collagenase solution was thawed and pre-heated at 37°C for 30 min. Samples were pre-incubated in 1X HBSS buffer at 37°C for 10 min before being moved to pre-heated collagenase solution at 37°C. Permeabilization was followed by a post-fixation step using 4% PFA for 20 min and then washed in 1 X PBSN (1xPBS with 0.1% sodium azide). Samples were incubated overnight in PBST (1% Triton) and 5% Normal Donkey Serum (NDS). Primary antibodies were incubated with the samples for 2 days in PBST and 5% NDS. The antibodies list is available in Supplemental Table1. For isotype controls, non-specific Goat IgG (02-6202, ThermoFisher Scientific), was incubated with samples for 2 days in PBST and 5% NDS. Primaries antibodies were washed in PBST overnight. Samples were incubated with conjugated secondary antibodies, at a concentration of 0.8 µg/µl, for 2 days in secondary sample buffer (LifeCanvas Technologies, USA) and washed in 1X PBSN overnight. For secondary controls, Donkey anti-Goat IgG secondary antibodies were incubated with samples without prior incubation with primary antibodies. Oxazole Yellow, (YO-PRO1, 40089, Biotium) was added to the secondary antibody solution with a dilution of 1:1000 to label nuclei (final concentration of 1 μM). Secondaries were washed overnight in PBSN. Samples were then fixed in 4% PFA for 20 min.

In this work we used traditional antibodies (Immunoglobulin G) to label various proteins such as Sclerostin, Connexin 43 (CNX43), Osteopontin (OPN), Osteocalcin (OCN), and Matrix metalloproteinase 9 (MMP9). Description of those targeted proteins and the list of antibodies used can be found in Supplemental Table 1.

### Whole-mount immunofluorescence, via nanobodies

Following decalcification, samples were preserved using SHIELD (LifeCanvas Technologies, USA). Contrary to the antibody method described above, the enzymatic permeabilization step is not required in this approach due to the smaller size of the nanobodies compared to antibodies. Indeed, nanobodies, or single variable domain on a heavy chain (VHH), are produced by camelids. They are relatively small molecules (∼ 15kDa) that correspond to the antigen-binding fragment of a heavy chain. Their size (∼10 times smaller than IgG) makes them particularly interesting in the context of diffusion in dense extracellular matrix. If the bone marrow is left in the tissue, a clearing step is needed to remove lipids. Samples were incubated in Clear+ Delipidation buffer (LifeCanvas Technologies, USA) for 3 days at 45°C. In addition, to remove the remaining non-protein elements, we adapted the F-Disco method [15], [30]. F-Disco is a tissue-clearing approach using organic solvents that has been shown to effectively clear highly heterogenous tissues. Samples were dehydrated with tetrahydrofuran (THF) through an increasing graded series of incubations (50%, 70%, 80%, 95%, 95% of THF) diluted in 75% of 1xPBS and 25% Quadrol (PBSQ). Quadrol is thought to reduce tissue autofluorescence [31]. For each step, samples were incubated for 30 min at 4°C. Once dehydrated samples were incubated in 100% dichloromethane (DCM) for 36 hours at 4°C with shaking, with several changes of DCM. DCM creates inverse lipid micelles allowing lipid removal. Samples were then rehydrated by decreasing the concentration of THF (95%, 95%, 80%, 70%, 50% of THF) in PBSQ, 30 min per step at 4°C. Samples were washed in water with several replacements for 24 hours at 4°C. Autofluorescence of the bone marrow and external tissues was further reduced by incubating samples in 3% hydrogen peroxide for 3h in a custom photobleaching box equipped with LED lights.

As a blocking step, samples were incubated overnight in PBS with 1% Triton-X (PBST) and 5% NDS. Anti-sclerostin nanobodies were custom-made by Creative Biolabs and conjugated with Alexa-Fluor 647 (TAB-388CT, Creative Biolabs). Nanobody solution was prepared in PBST with 5% NDS and incubated for 2 days with the samples. Bones were washed in PBST overnight. The signal was amplified using anti-camelid-nanobodies fragment antigen-binging region (Fab), conjugated with AF647, for 2 days in secondary sample buffer (LifeCanvas Technologies, USA). Oxazole Yellow (1:1000) was added to the secondary antibodies for nuclear staining. Secondaries were washed in 1x PBSN overnight.

### Refractive index matching and sample mounting

Refractive index matching is a critical step to enable light to penetrate through the samples during imaging while limiting scattering. To do so, EasyIndex (LifeCanvas Technologies, USA) was used. EasyIndex has a refractive index of 1.52, similar to collagen’s refractive index. Samples were incubated in 100% EasyIndex overnight with shaking. Mounting gel was prepared by mixing EasyIndex and ultra-low melt agarose. The final concentration of agarose was 2%. The gel was heated to 90°C and cooled down to enable mounting. Gel was poured in a 3D printed holder and long bones were mounted longitudinally with anterior part of the tibia facing up. Detailed mounting protocols can be found in the supplemental methods section.

### Lightsheet microscopy

Whole-mount fluorescent images were acquired with LifeCanvas Technologies’ SmartSPIM (lasers wavelengths - emission filters: 488nm - Chroma ET525/50m, 561 nm - Chroma ET600/50m, and 647 nm - Chroma ET690/50m). The step size along the Z-axis was 2 µm. Samples were imaged using a 3.6x objective (1.8 µm x 1.8 µm x 2 µm voxel size) or 9x objective (0.7 µm x 0.7 µm x 2 µm voxel size). During imaging, samples were immersed in EasyIndex matched immersion oil RI=1.52 (LifeCanvas Technologies, USA). A 16-bit greyscale Tiff stack was acquired for each channel. Tile scans were acquired with 10% overlap along the longitudinal axis of the bone. A stack of 16-bit greyscale TIFF images was acquired for each channel. Image stacks were stitched and de-stripped using the SmartSPIM post-procession pipeline (LifeCanvas Technologies). TIFF images were converted to Imaris files for 3D visualization. 3D reconstruction of the lightsheet images were generated using Imaris 9.5.1 (Oxford Instrument, UK).

### Confocal microscopy

3D index-matched samples were placed on a coverslip with a drop of EasyIndex. On the other side, the coverslip was customized to hold water for objective immersion. Samples were imaged using the Leica TCS SPE (Leica, Germany) with a 20x water objective (HCX APO L U-V-1, NA: 0.5), laser wavelengths/filters of 488 nm/500-590nm, 561nm/570-627nm, and 635 nm/650-750nm, a gain of 800 V, and a frame average of 2.

### Cell detection

A custom convolution neural network was used for automated cell detection (LifeCanvas Technologies, USA). Two networks in sequence were involved. First, possible cell locations were found via a fully-convolutional detection network [32] based on a U-Net architecture [33]. This network is used for semantic segmentation and was trained to generate candidate cell locations while minimizing the number of false negatives. Second, each possible cell location was classified as either positive or negative via a convolutional network using a ResNet architecture [34]. This classifier filters false positives generated during the segmentation step, resulting in more precise cell detection.

The main output is the centroid of each cell. MATLAB (MathWorks, USA) and Napari [35] were used to visualize cell centroids as individual points and to quantify the number of detected cells (Figure 2). External tissues and remaining bone marrow pieces were segmented out in ImageJ to isolate the cortical bone. The mask was applied to the cell detection data to include only cells within the cortical bone, resulting in a masked cell detection point cloud (Figure 2.C). To normalize the sample length, midshafts were isolated starting from 1.5 mm proximal to the proximal tibia crest to the distal tibia-fibula junction. Tibia midshafts were discretized in 20 µm thick sections, and the number of cells detected in these sections was reported along the midshaft length (Figure 2.C). The cell detection pipeline is summarized in Figure 2.

**Figure 2:**
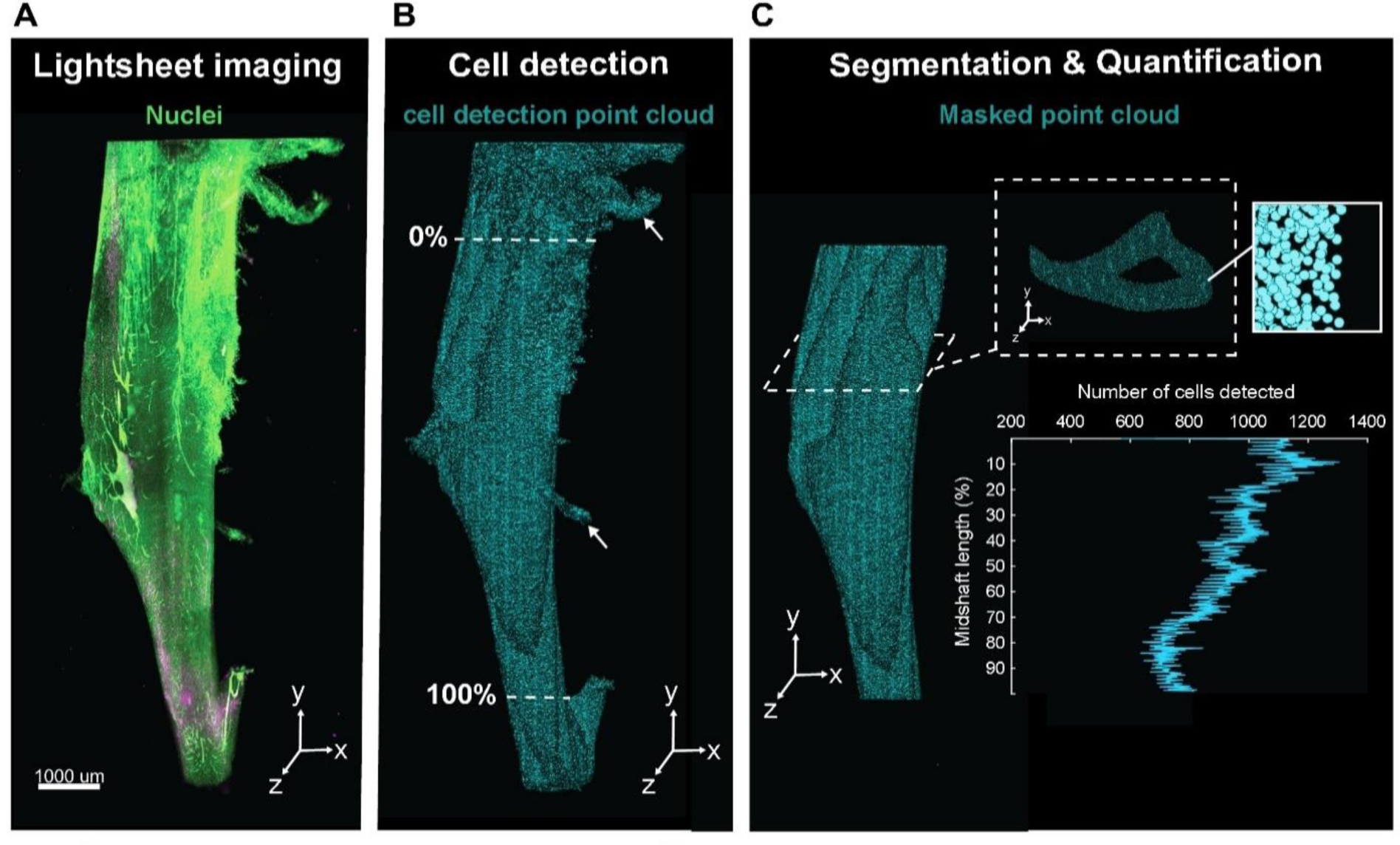
Summary of the images’ quantification pipeline. A) Lightsheet images were used as input in a custom cell detection model. B) Cell detection resulted in a list of points corresponding to individual centroid of detected cells. Together all centroids form a point cloud. Raw cell detection point cloud showed detection in external tissues pointed by white arrows. C) Samples were manually segmented to isolate the cortical bone and remove unspecific detection, resulting in a masked point cloud. The number of cells detected within 20 µm-thick sections were reported along the normalized midshaft length.

## Results

### Whole-mount mRNA labeling

We labeled and imaged *Sost* and *Dmp1* mRNA. These two genes have been shown to play an important role in the regulation of bone adaptation and homeostasis. *Sost* encodes the protein sclerostin, which inhibits bone formation and is expressed by mature osteocytes [36]. *Dmp1*, expressed in late osteoblasts and osteocytes, is critical for bone mineralization [37], [38]. A *Dmp1* promoter is also commonly used to drive expression of endogenous fluorescence in osteocytes [39]. All mRNA labeling was performed in the mouse tibia midshaft. The midshaft is the bone region where most of cortical adaptation occurs in the murine tibial bone loading model [40], [41].

Figures 3.A shows a mouse tibia 3D reconstruction of the cells expressing mRNA for *Sost*. Virtual re-sectioning was performed to show examples of cross-sectional 2D slices showing osteocytes expressing *Sost* (Figure 3.B). Figures 3.C shows a longitudinal 2D view of the cortical bone where *Sost* and *Dmp1* mRNA are expressed. Signal colocalization is also accessible, as shown in Figure 3.C and D, where *Dmp1* and *Sost* signal are overlapping (indicated by white cells in Figure 3.C). In Figure 3.D, there are overlaps between osteocytes expressing *Sost* mRNA and osteocytes nuclear staining. In comparison, no signal was observed in osteocytes for the negative control conditions (Figure 3.E).

**Figure 3:**
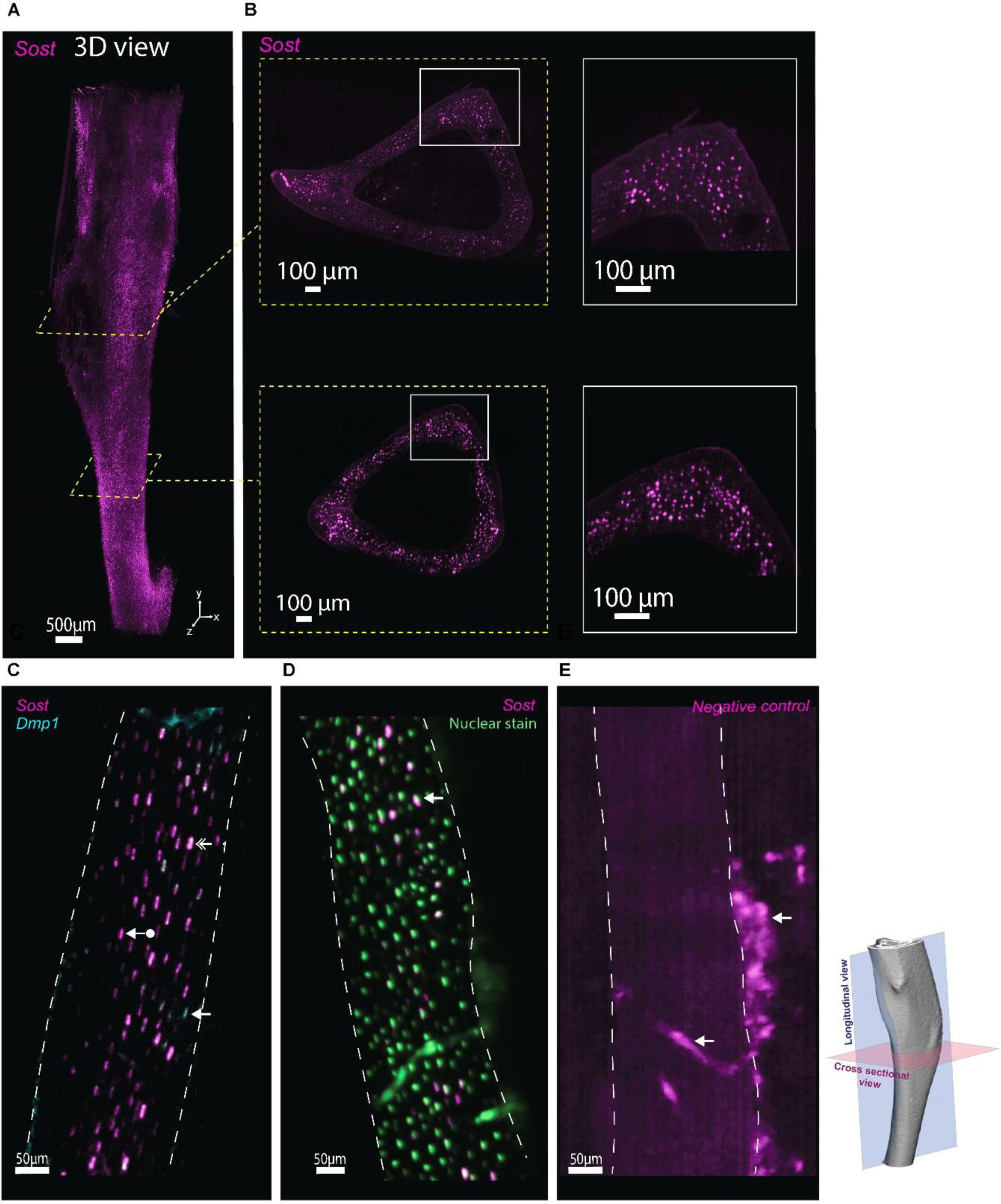
Whole mount HCR FISH in mouse tibia. A) 3D view of a mouse tibia midshaft labeled for Sost. B) Cross-sectional 2D view of the 3D mouse tibia midshaft showing Sost expression in the cortical bone. C) Zoom on the cortical bone showing expression of Sost (arrow with circle) and Dmp1 (plain arrow) through the cortical thickness. Osteocytes expressing both Dmp1 and Sost appear white (double-headed arrow). D) Zoom on the cortical bone showing colocalization of Sost signal and nuclear staining (plain arrow). E) Negative control. Sample was incubated with fluorescent DNA hairpins without prior incubation with probes. White arrows show autofluorescence of a blood vessel and some remaining part of bone marrow. Images were acquired via lightsheet microscopy. Dotted lines highlight the cortical bone.

In this work, we were interested in labeling and imaging osteocytes located in the cortical bone. Figure 4.A is a masked point cloud of all the detected cells expressing *Sost* mRNA in the cortical bone (Figure 4.A). The number of cells expressing a gene of interest can then be normalized by the total number of cells (nuclear staining detection). In the cortical bone of the midshaft, we detected, on average, more than 400,000 cells (n=3) based on nuclear staining. On average, 65% of the cells were expressing *Sost* which corresponds to the previously reported range of osteocytes expressing sclerostin in control mouse bone [42], [43], [44]. Figure 4.C plots the percentage of cells expressing *Sost* along the tibia midshaft length.

**Figure 4:**
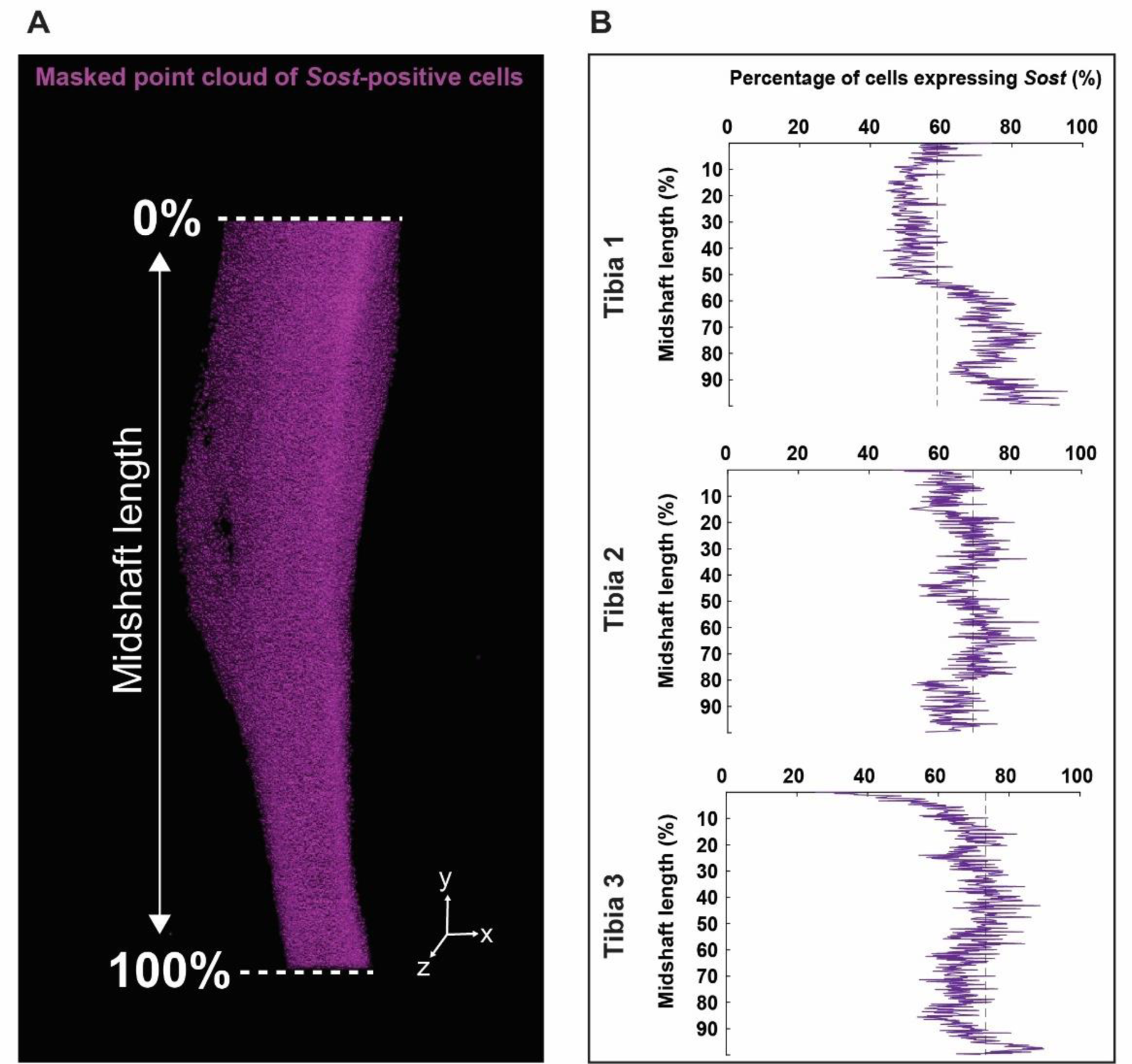
Whole-mount cell detection in mouse tibia. A) Masked cell detection point clouds including only cells in cortical bone expressing Sost mRNA. B) Percentage of cells expressing Sost along the bone length of three different samples. Each data point corresponds to the number of cells expressing Sost mRNA in osteocytes in a 20 µm thick section.

### Whole-mount immunofluorescence via antibodies

Osteocytes are embedded in a dense extracellular matrix mostly made of collagen. Penetration of antibodies in bone is challenging due to their size (∼150 kDa), contrasting with DNA oligos used in HCR-FISH mRNA labeling, which are much smaller than antibodies. In this method, we used a matrix permeabilization step to facilitate the diffusion of antibodies in the decalcified bone. To permeabilize the matrix and prevent sample degradation, bones were preserved using SHIELD (LifeCanvas Technologies, USA) [11]. After isolation of the tibia midshaft, samples were permeabilized using collagenase. Figure 5 shows that the overall bone structure was maintained after the permeabilization step.

**Figure 5:**
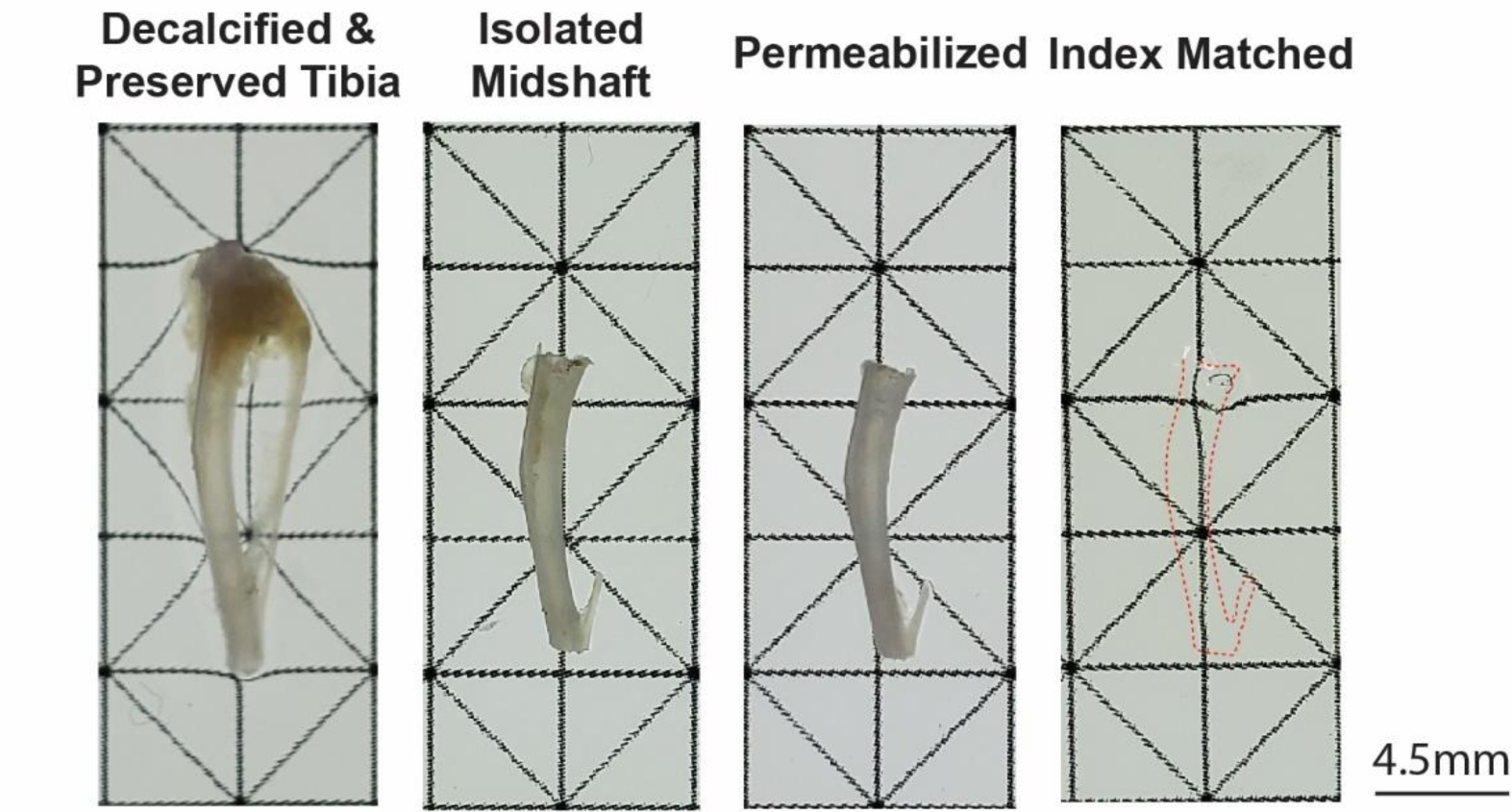
Mouse tibia images taken after each step of the method. After the tibia was collected, the bone marrow was flushed. The sample was then fixed, decalcified, and preserved using SHIELD. The tibia midshaft was then isolated and permeabilized using collagenase. After labeling, the sample was index-matched to RI= 1.52. The red dotted line shows the contour of the index-matched sample.

We found that delipidation was not required to achieve full optical transparency due to the initial removal of bone marrow and external tissues from the tibia midshaft. After permeabilization, incubation in EasyIndex resulted in optical transparency (Figure 5)

We found that the permeabilization step was critical for the complete penetration of antibodies. Figure 6 shows the effect of collagenase permeabilization on the diffusion of antibodies through cortical bone. One hour of permeabilization led to only surface penetration. Longer permeabilization times allowed the antibodies to diffuse through the cortex (Figure 6), but over-permeabilizing degraded the structure (Supplemental Figure 5). Optimization of permeabilization timing is needed, based upon the density of collagen, the thoroughness of demineralization, and the potency of the collagenase batch (which varied from batch to batch). We found that SHIELD helped achieve matrix permeabilization while limiting tissue damage. In Supplementary Figure 6, we compared the morphology of SHIELD-preserved and non-preserved femur midshafts after three hours of permeabilization. The SHIELD-preserved samples maintained their morphology whereas the non-preserved samples were almost fully digested. The addition of 1% SHIELD Epoxy Solution in SHIELD ON Buffer protected sample edges from over-permeabilization while allowing full penetration of the antibodies after 6h of permeabilization (Supplemental Figure 4). Antibody penetration occurred from both endosteal and periosteal surfaces, as well as the vascular canals (Supplemental Figure 3). Optimal permeabilization time was found to vary between samples. For a tibia from an adult mouse, without external tissue or bone marrow, and treated as described above, we found that 6h of collagenase permeabilization showed optimal antibody penetration. Trials are advised to determine optimal timing of permeabilization. Although permeabilization was found to be necessary for antibody penetration, permeabilization is not required to achieve optical transparency of the sample.

**Figure 6:**
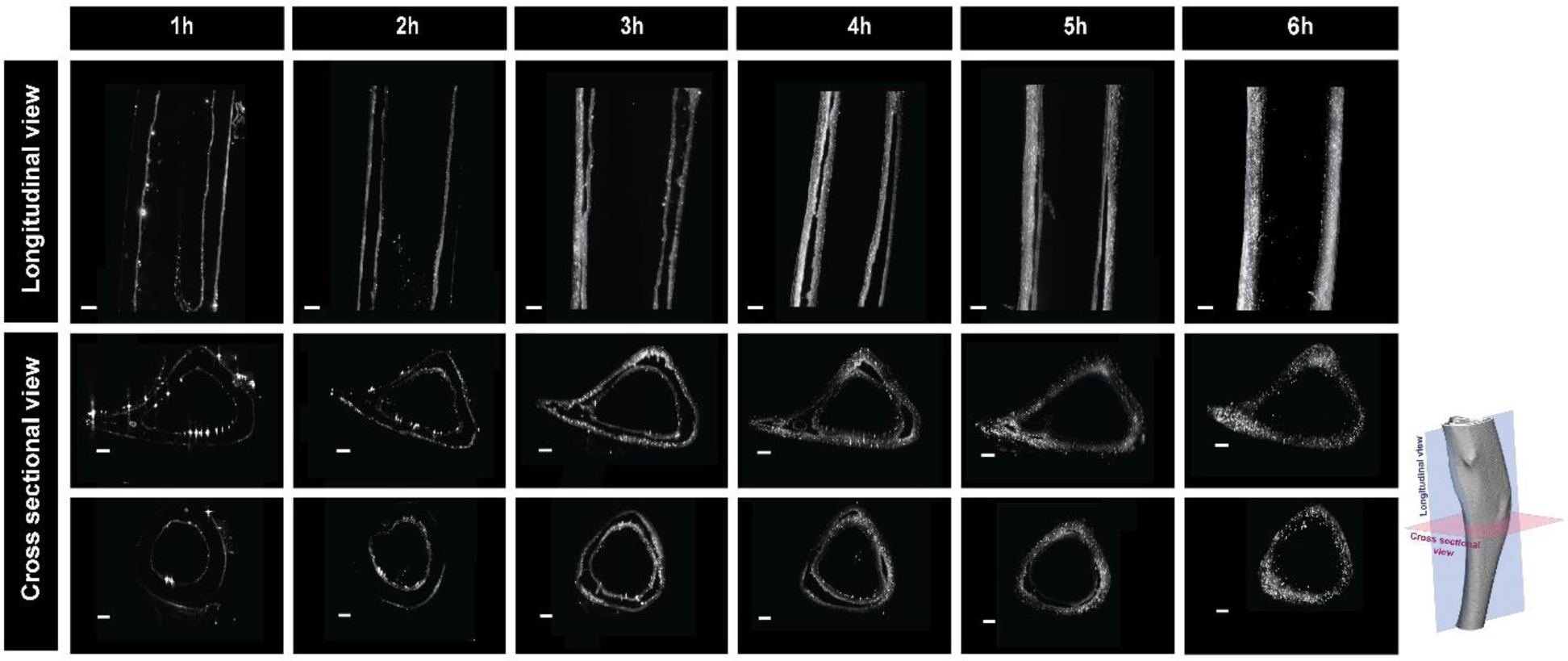
Effect of collagenase permeabilization on antibody penetration (Anti-Sclerostin) through the cortical thickness of SHIELD-preserved samples. Longitudinal view of mouse tibiae. Short permeabilization steps led to surface penetration from both periosteal and endosteal surfaces. Longer permeabilization times enabled full penetration of the antibodies throughout the cortical thickness. Scale bar corresponds to 150 µm.

Using WISH-BONE, we labeled sclerostin in 3D mouse tibia midshafts (Figure 7.A) throughout the cortical bone (Figure 7.B, 7.C, 7.D). Sclerostin is a protein highly expressed by mature osteocytes and is known to inhibit the bone adaptation process. Sclerostin has been reported to be downregulated in loaded mouse legs compared to control legs [43]. Regulation of sclerostin in mechanoadpatation studies is often reported via 2D immunolabeling, and sclerostin-positive cells are manually counted in a region of interest [43], [45].

**Figure 7:**
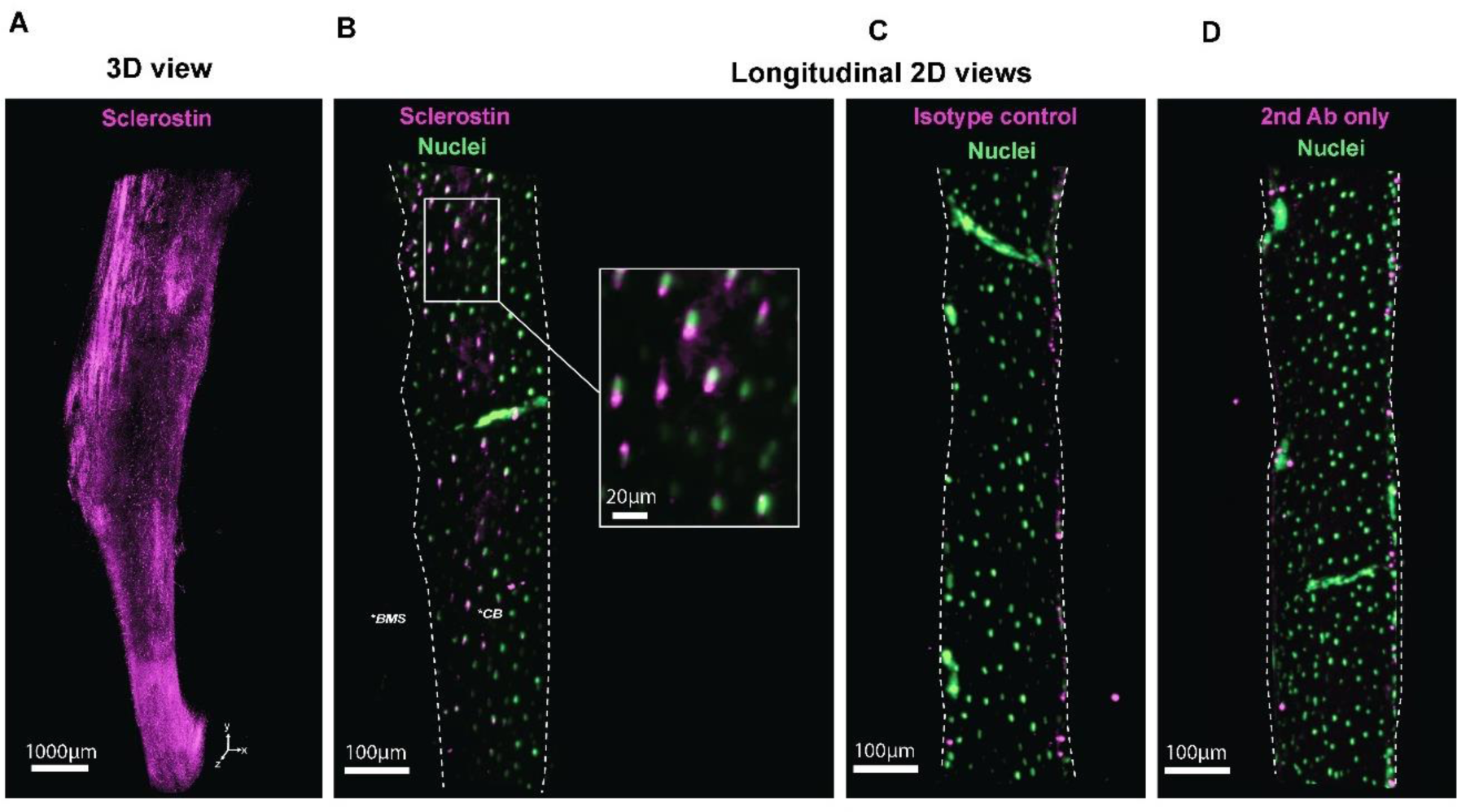
Whole-mount immunolabeling of sclerostin in mouse tibiae. A) 3D view of a mouse tibia midshaft labeled for sclerostin acquired with lighsheet microscopy. B) Longitudinal 2D view of the 3D mouse tibia midshaft showing sclerostin expression throughout the cortical thickness. C & D) Longitudinal 2D views of 3D mouse tibia midshafts, respectively, showing the results of the isotype control and secondary antibody control.

In this work, signal from the anti-Sclerostin Goat IgG can be observed through the cortical thickness (Figure 7.B). This signal was lacking in isotype and secondary controls (Figure 7.C, 7.D). In the isotype control condition Goat primary antibodies lacking target specificity were used. Isotypes control confirmed the specificity of the primary antibody used to target sclerostin (Figure 7.B). Secondary controls used only fluorescently labeled secondary antibodies omitting the use of primary antibodies to assess the potential non-specific binding of the secondary antibodies. Together, these controls validate the signal observed in osteocytes in Figure 7.B.

Osteocytes are located in lacunae which are connected to one another via canaliculi [46]. Connexin 43 (CNX43) are gap junctions that have been shown to connect osteocytes’ dendrites and allow cell-to-cell communication [47] and play an important role in bone homeostasis [48]. Disruption of this cell-to-cell communication could lead to a decrease in bone mechanosensitivity.

Figure 8.A shows a 3D reconstruction of a mouse tibia labeled for CNX43. CNX43 signal can be observed around each osteocyte nucleus showing where osteocyte dendrites connect with their neighbors. Signals were observed in both cortical (Figure 8.B) and trabecular bone (Figure 8.C). Sample clearing and imaging resolution allowed us to visualize gap junctions as dots around the osteocytes’ nuclei (Figure 8.B)

**Figure 8:**
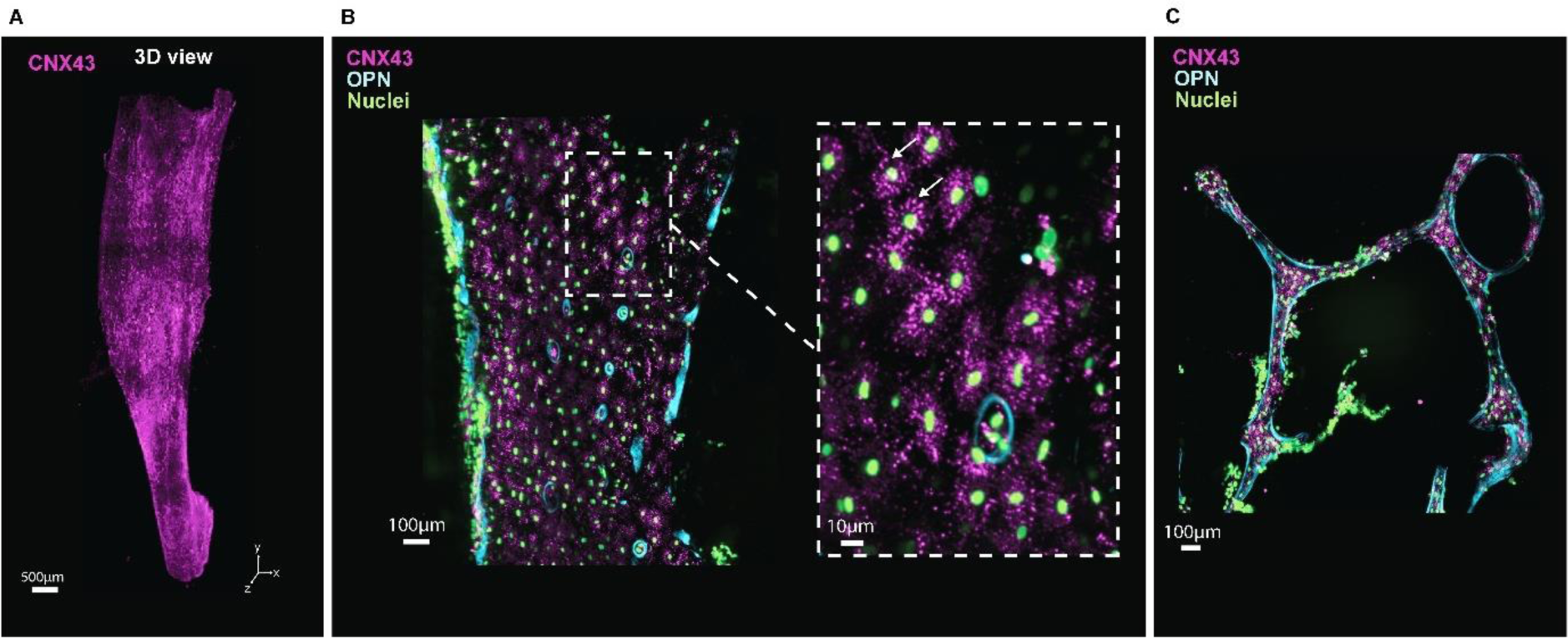
Whole-mount immunolabeling of CNX43 and OPN in a mouse tibia. A) 3D view of a mouse tibia midshaft labeled for CNX43 acquired with lighsheet microscopy. B) Longitudinal 2D view of the 3D mouse tibia midshaft showing CNX43 and OPN expression in the cortical bone. Zoom on the cortical bone showing CNX43 expression around the osteocytes (white arrows). C) 2D view of the 3D mouse tibia midshaft trabecular bone showing CNX43 and OPN. Images were acquired at 9x via lightsheet microscopy.

In this work, we also labeled and imaged osteopontin, which plays an important role in bone cell adhesion and migration by binding to cell surface integrins via an RGB sequence. It has been shown to be implicated in bone metabolism and in bone-related diseases such as osteoporosis or osteosarcoma [49]. OPN signal was observed at the endosteal and periosteal surfaces of the cortical bone, where most of the osteoclast activity is expected (Figure 8.B, 10.A, 10.B). We also imaged OPN signal along trabeculae left in the proximal end of the midshaft (Figure 8.C, 10.A, 10.B).

Osteocalcin is the most abundant non-collagenous protein in the bone and is expected to be present throughout the entire bone thickness. It is secreted by osteoblasts and plays a role in bone mineralization. OCN has been suggested to form a bridge between mineral and bone matrix by binding to hydroxyapatite and forming complexes with collagen [50]. Following preservation, permeabilization and antibody staining, OCN signal was observed throughout the cortical bone. Figure 8.A shows a longitudinal 2D virtual slice of cortical bone labeled for OCN and MMP9. OCN staining highlights the lamellar structure at the periosteal surface whereas a more woven bone structure is observed deep in the cortical bone. Using Imaris (Oxford Instrument, UK), images were virtually re-sectioned to access cross-sectional 2D views of the samples (Figure 9.B & 9C). Virtual cores-sectional views show the difference in the structural organization at the periosteal surface compared to the endosteal surface. It is important to note that virtually re-sectioning in a different plane will result in a loss of resolution due to the anisotropy of the voxel size.

**Figure 9:**
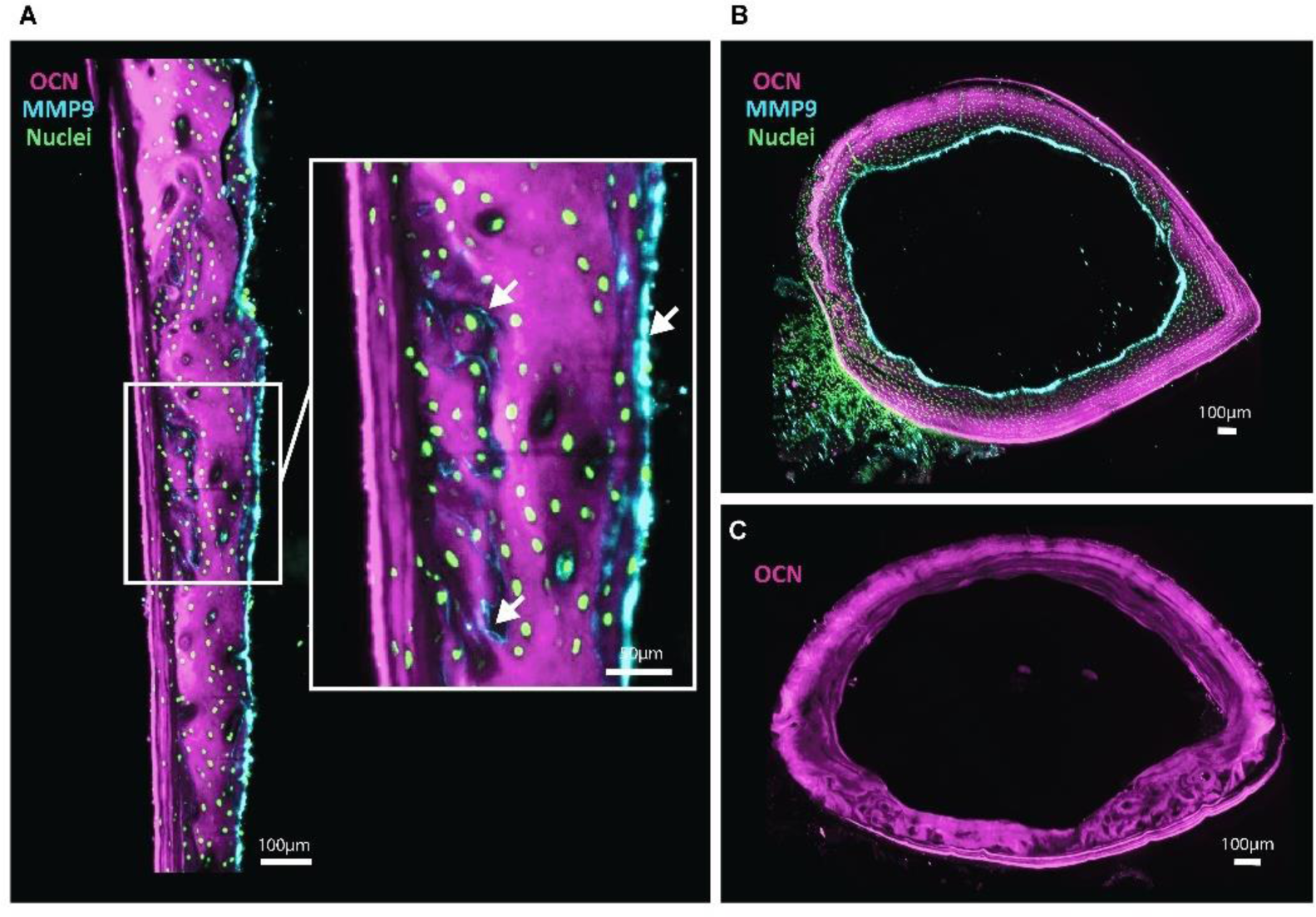
Whole-mount immunolabeling of osteocalcin (OCN) and matrix metalloproteinase-9 (MMP9) in a mouse femur. A) Longitudinal 2D view of the 3D mouse tibia midshaft showing OCN and MMP9 in the cortical bone. The inset shows a zoom in the cortical bone. White arrows point to regions of MMP9 expression. B & C) 2D cross-sectional view of the 3D mouse tibia midshaft. Images were acquired at 9x via lightsheet microscopy.

Using the same pipeline, we observed that MMP9 signal was mainly observed at the bone surfaces. Some thin lines of MMP9 expression were observed within the woven bone matrix (Figure 9.B). MMP9 is a metalloproteinase expressed by mature osteoclasts [51]. It plays an important role in matrix degradation and activation of growth factors. MMP9 has also been shown to participate in the migration of preosteoclasts [52].

Using confocal microscopy, we acquired specific locations of the 3D labeled samples, with higher image resolution compared to lightsheet microscopy. Due to the transparency of the samples, confocal images could be obtained from the middle of the tissues. Similar to lightsheet microscopy, we observed OCN expression in a trabecula (Figure 10.B) and CNX43 signal around osteocytes in cortical and trabecular bone (Figure 10A & 10.B).

**Figure 10:**
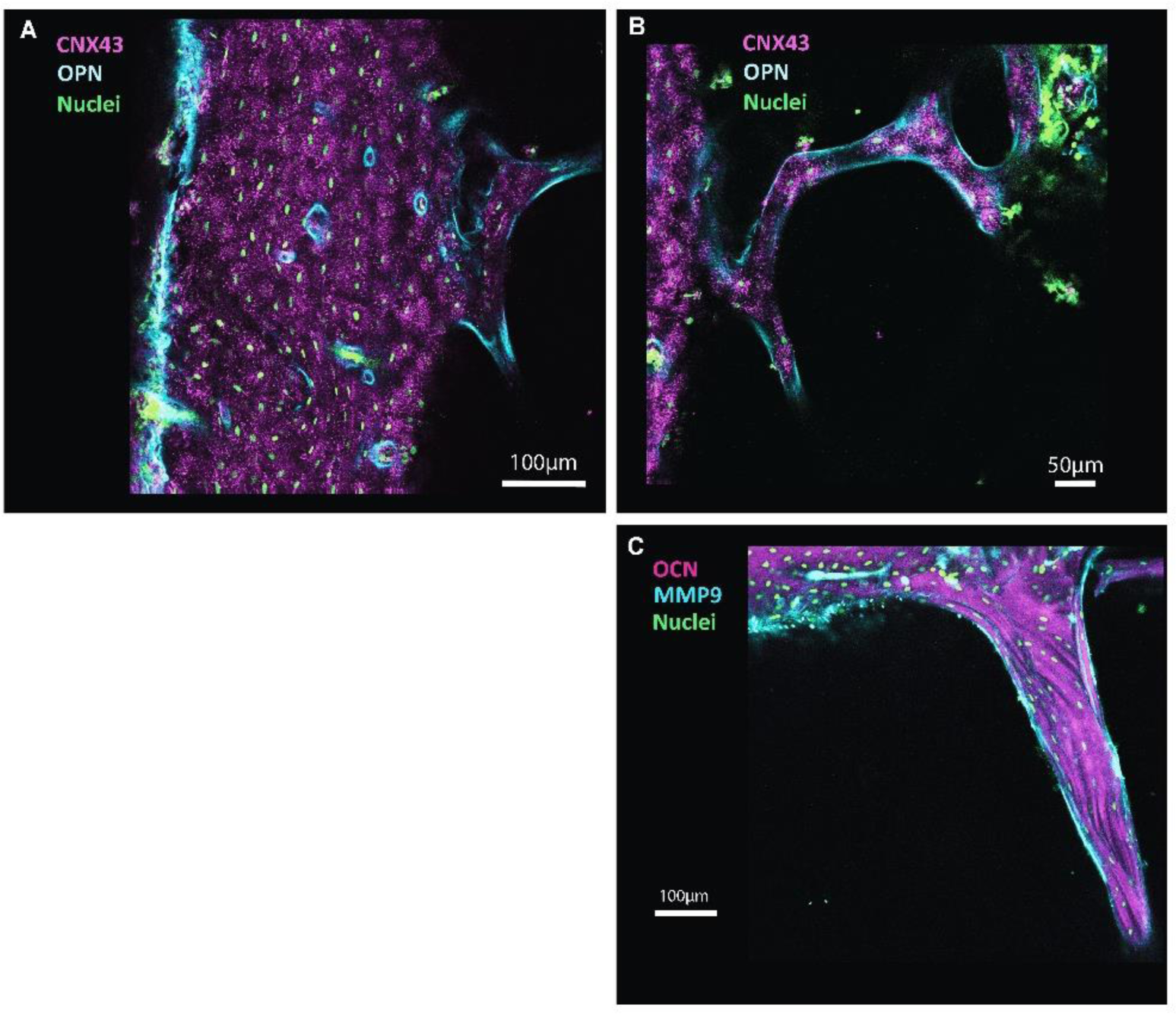
Confocal images of 3D cleared and labeled mouse bone samples. A) & B) 2D longitudinal view of the 3D mouse tibia midshaft showing CNX43 and OPN expression in cortical and trabecular bone. C) 2D longitudinal view of the 3D mouse bone sample showing OCN and MMP9 expression in a trabecula. Images were acquired at 20x via confocal microscopy.

### Whole-mount immunofluorescence via nanobodies

Nanobodies are particularly promising in the context of diffusion through a dense collagen matrix due to their relatively small molecular weight. In this study, we investigated the use of customized camelid VHH to label sclerostin in 3D mouse bone.

First, nanobodies were validated on 2D sections of adult mouse femurs (Supplemental Figure 2). Confocal images showed that nanobodies, specific to sclerostin, labeled osteocytes. No signal in osteocytes was observed in negative control conditions which only involved secondary antibodies.

Next, we acquired signals from nanobodies in whole-mount mouse femurs and vertebrae (Figure 11 & 12). As VHHs do not require matrix permeabilization, bone marrow and external tissues were kept intact. However, the samples were delipidated as the lipid in the bone marrow would result in large light scattering. (Delipidation can be avoided by flushing the bone marrow out and removing all external tissues). Figure 11 shows the 3D reconstruction of a mouse femur; the green channel was used to visualize autofluorescence, and the far-red channel was used to label sclerostin. As described previously, multiple 2D views of the sample can be accessed to investigate specific regions of the sample (Figure11.B to 11.E). Figures 11.C & 11.D focus on the femoral neck and femoral head. We observed a high background signal in the far-red channel (Figure 11.C) compared to endogenous autofluorescence (Figure 11.D). However, zooming in to the cortical bone showed fluorescence from sclerostin labeling in osteocytes.

**Figure 11:**
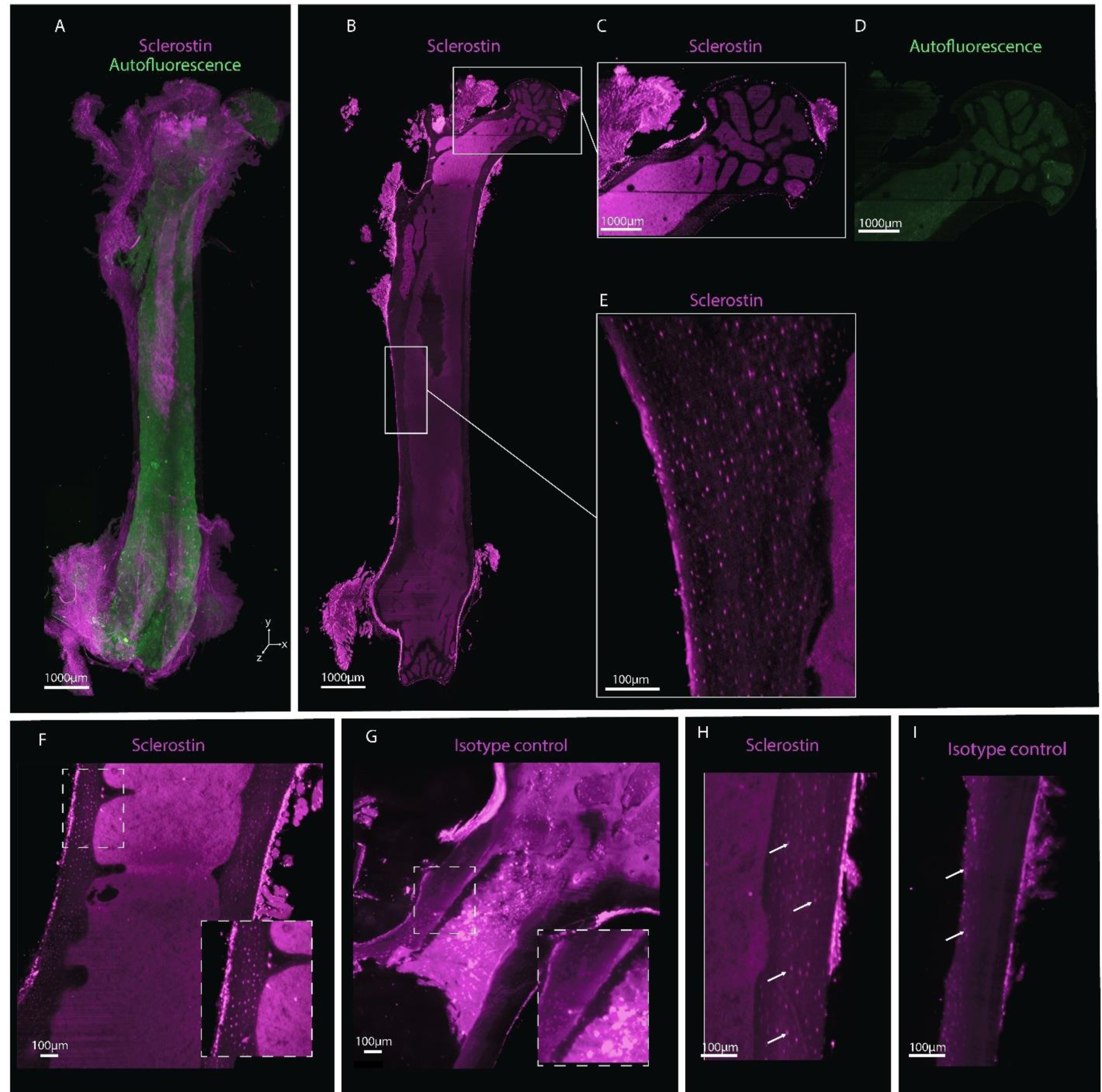
Whole-mount immunolabeling of a whole mouse femur using anti-sclerostin nanobodies. A) 3D view of a cleared mouse femur. B) Longitudinal 2D view of a 3D femur. C) Zoom on the femoral head. D) Autofluorescence. E) Zoom on the cortical bone showing osteocytes expressing sclerostin. F) Femoral neck labeled with anti-sclerostin nanobodies. G) Femoral neck labeled with anti-rabbit-IgG nanobodies. H) Femoral midshaft showing osteocytes expressing sclerostin (white arrows). I) Femoral midshaft showing non-specific signal from osteocytes (White arrows).

The nanobodies’ signal was relatively dim even with Fab fragment amplification in whole-mount femur, which could limit the use of nanobodies. Lightsheet images could not confirm signal from nanobodies in the trabecular bone due to high background signal.

The signal from anti-sclerostin nanobodies was also imaged in mouse vertebra. Figure 12.A displays a cross-sectional 2D view showing nuclear staining, autofluorescence, and signal from anti-sclerostin nanobodies. The combination of delipidation buffer (LifeCanvas Technologies, USA) and fDISCO [15], [30] resulted in cleared samples despite the external tissue and bone marrow (Figure 12.A & 12.B). Sclerostin signal was observed in cortical bone (Figure 12.E & 12.E’). As isotype control, a vertebra was incubated with nanobodies targeting rabbit IgG. Figure 13 shows the difference in signal obtained from anti-sclerostin nanobodies and isotype control in the spinous process. Sclerostin signal can be observed in the osteocytes of the samples stained with anti-sclerostin VHH, whereas the signal is lacking in the isotype control.

**Figure 12:**
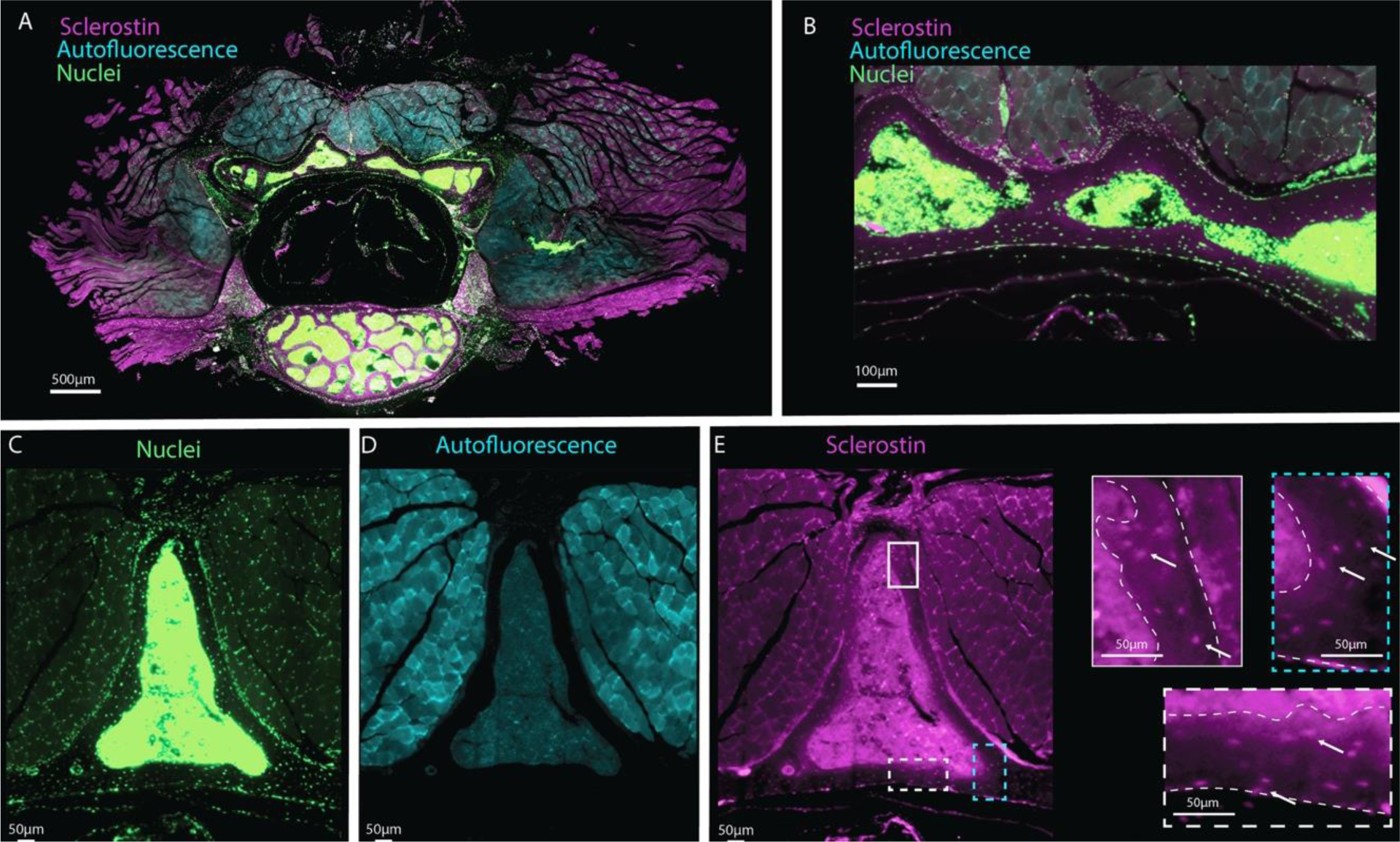
Whole mount immunolabeling of a mouse vertebra using anti-sclerostin nanobodies . A) 2D view of 3D cleared mouse vertebra. B) Zoom on a region showing cortical bone, osteocytes nucleus, bone marrow, and external tissue. C-D-E) 2D view of the neural spine respectively showing nuclei, autofluorescence, and anti-sclerostin nanobodies signals. E’) Autofluorescence. E) Zoom on the cortical bone showing osteocytes expressing sclerostin. Dashed lines highlight the cortical bone. All images were acquired using lightsheet microscopy.

**Figure 13:**
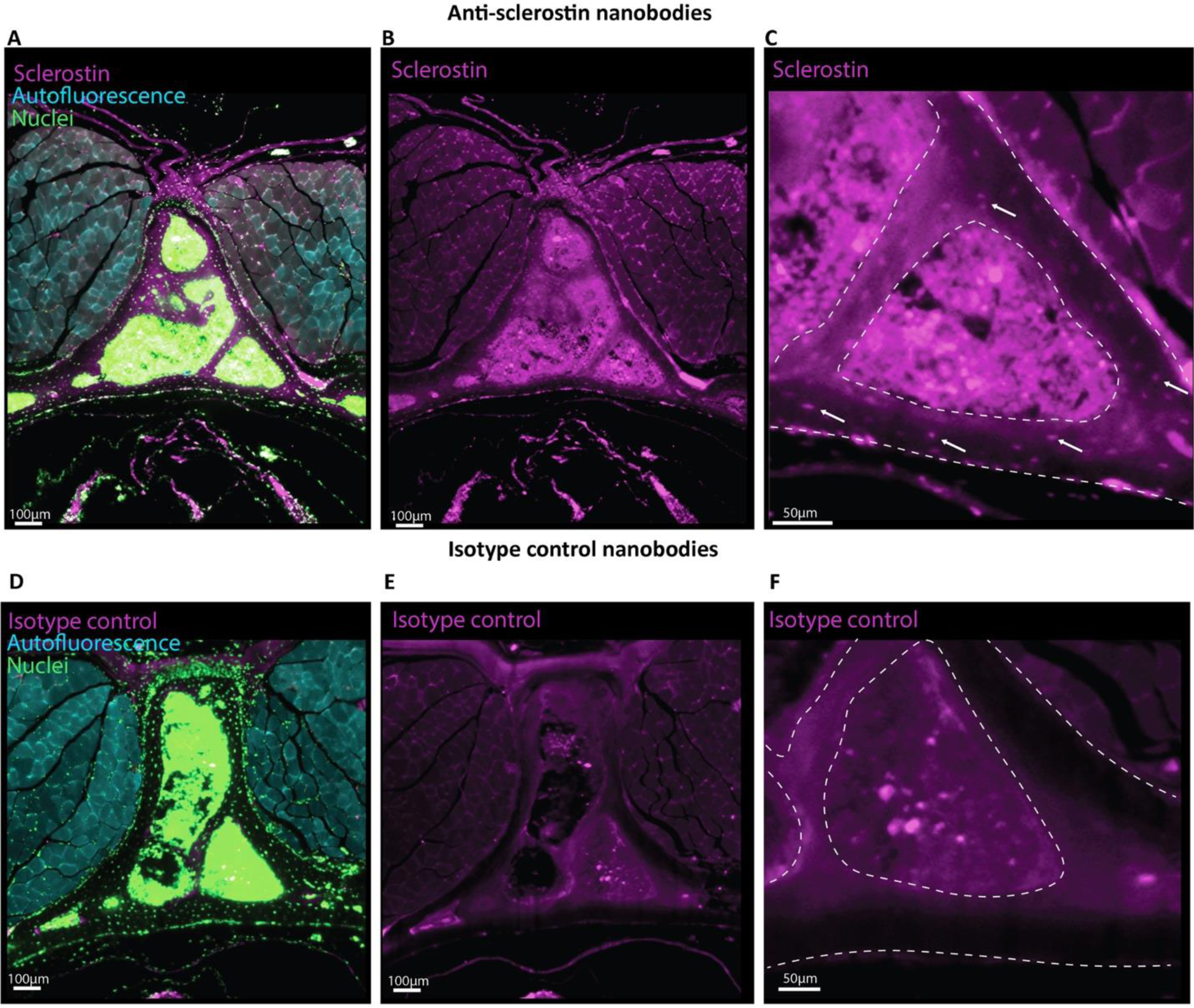
Comparison of sclerostin labeling and isotype control in mouse vertebra using nanobodies. A-B) 2D view of 3D cleared mouse vertebra labeled with anti-sclerostin nanobodies (A, B, C) and nuclear stain (A). C) Zoom on the neural spine region showing cortical and trabecular bone, and sclerostin-positive osteocytes. D-E) 2D view of 3D cleared mouse vertebra labeled with isotype control (anti-Rabbit-IgG nanobides) (D, E, F) and nuclear stain (D). C) Zoom on the neural spine region showing cortical, and trabecular bone. Dashed lines highlight the cortical bone. All images were acquired using lightsheet microscopy.

## Discussion

We described WISH-BONE, a method enabling osteocyte molecular labeling in the 3D mouse bone. This approach addresses the limitation of current methods for mRNA and protein analysis in bone by providing three-dimensional information on the biomolecule expression. We showed that mRNA and protein labeling can be achieved throughout the cortical bone. Cell detection enables quantification of the number of cells expressing a molecule of interest, offering new ways to investigate bone and osteocyte biology.

In this work, we showed that mRNA could be labelled as well as various protein types such as hemi-channels (Connexin 43), structural proteins (Osteocalcin), secreted glycoprotein (Sclerostin), and phosphorotein (Osteopontin), suggesting that this method could be used in various bone-related fields.

We plan on using this method to investigate bone mechanoadaptation. Our new method allows the investigation of hundreds of thousands of cells in the entire bone, providing a more complete interpretation of the impact of forces on the regulation of biomolecule expression.

Various parameters can affect outcome quality when using this method. We found that lot-to-lot variability in collagenase activity interfered with the time required in the permeabilization step. The collagenase used in this method had a specific activity between 125-250 Mandl units per milligram of powdered substance. Collagenase activity should be tested between lots to adjust permeabilization time. For an adult mouse tibia midshaft with removed external tissue and bone marrow, fully decalcified, and SHIELD-preserved, we recommend starting with 6 hours of collagenase permeabilization.

We found that shorter decalcification time could result in satisfying optical transparency and image quality. However, the level of decalcification might interfere with the permeabilization step. For immunolabeling using antibodies, we recommend thoroughly decalcifying samples. For mRNA labeling, decalcification time could be reduced as the DNA probes do not require tissue permeabilization to diffuse in the samples.

Epoxy concentration can be increased in the SHIELD solution to crosslink proteins more densely. This results in a longer permeabilization step to reach the full penetration of antibodies.

In this work, we showed that high-resolution images can be acquired even with external tissue and bone marrow left in the samples (Figure 11,12,13). This was made possible by the combination of aqueous and organic clearing approaches. Although, it does not decrease image quality, we noticed that bone marrow and external tissue might affect permeabilization of the cortical bone by collagenase. Thus, for immunolabeling using traditional IgG, we recommend flushing the bone marrow and dissecting out all external tissue to facilitate collagenase permeabilization and antibody penetration.

In addition, we recommend that any pre-treatment potentially affecting collagen stability, such as decolorization using hydrogen peroxide or delipidation using high temperature, be performed after the tissue has been permeabilized.

The combined use of epoxy preservation and enzymatic permeabilization provides a new approach to investigating bone biology. Our method provides 3D information of the osteocytes biomolecule expression which could be used to investigate the role of osteocytes in fracture healing, sarcoma, bone development, or mechanoadaptation.

## Conclusion

We presented new approaches to label and image mRNA and protein in 3D adult mouse bone. WISH-BONE allows visualization of osteocytes through the cortical bone thickness using epoxy preservation and collagenase permeabilization. This approach allows the investigation of hundreds of thousands of cells in the bone providing critical spatial information linking the molecular response and osteocyte 3D environment. Investigation of osteocyte molecular signaling is important in order to understand the regulation of the bone biology in the context of bone fracture healing, mechanoadaptation, development, and more. We anticipate this new tool to provide valuable insights into bone biology.

## Supporting information

Supplemental Results & Methods

## Acknowledgements

This work was funded by the National Science Foundation CMMI # 2010010 and the National Science Foundation INTERN supplement.

We thank the Institute for Chemical Imaging of Living Systems (RRID:SCR_022681) at Northeastern University for consultation and instrument support.

