## Supplemental Results & Methods for "WISH-BONE: Whole-mount In Situ Histology, to label osteocyte mRNA and protein in 3D adult mouse bones"

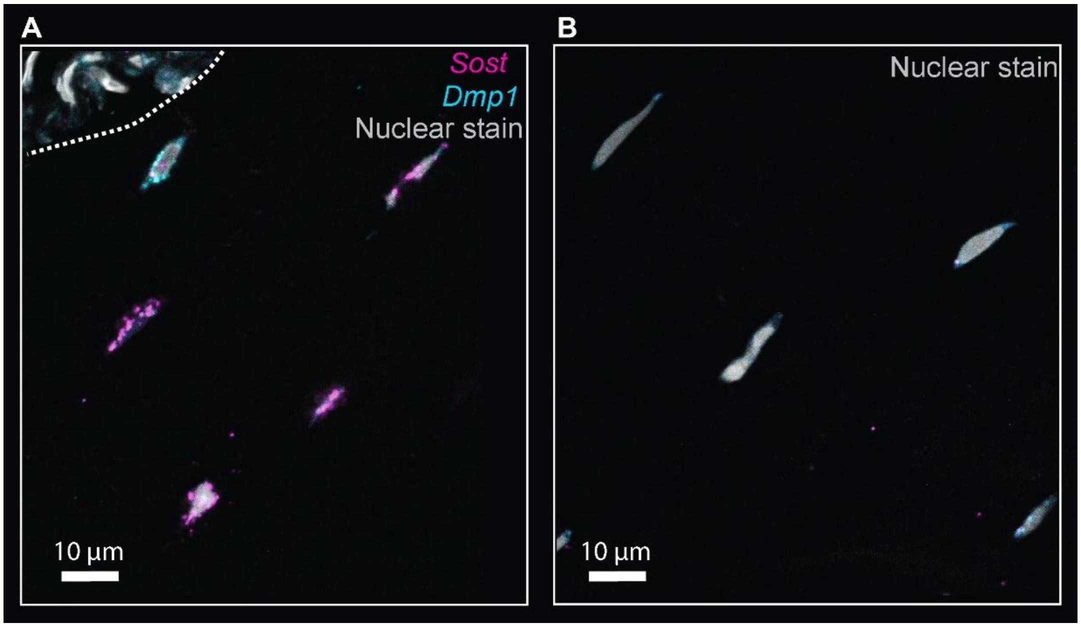

*Supplemental Figure 1: mRNA labeling via HCR FISH in mouse bone cryosections. A) Confocal images of osteocytes in cortical bone expressing Sost and Dmp1 mRNA. B) Confocal images of osteocytes showing autofluorescence of the tissue. Images were acquired with a 63x objective using Zeiss LSM 800, located in CILS facility at Northeastern University Boston, MA.*

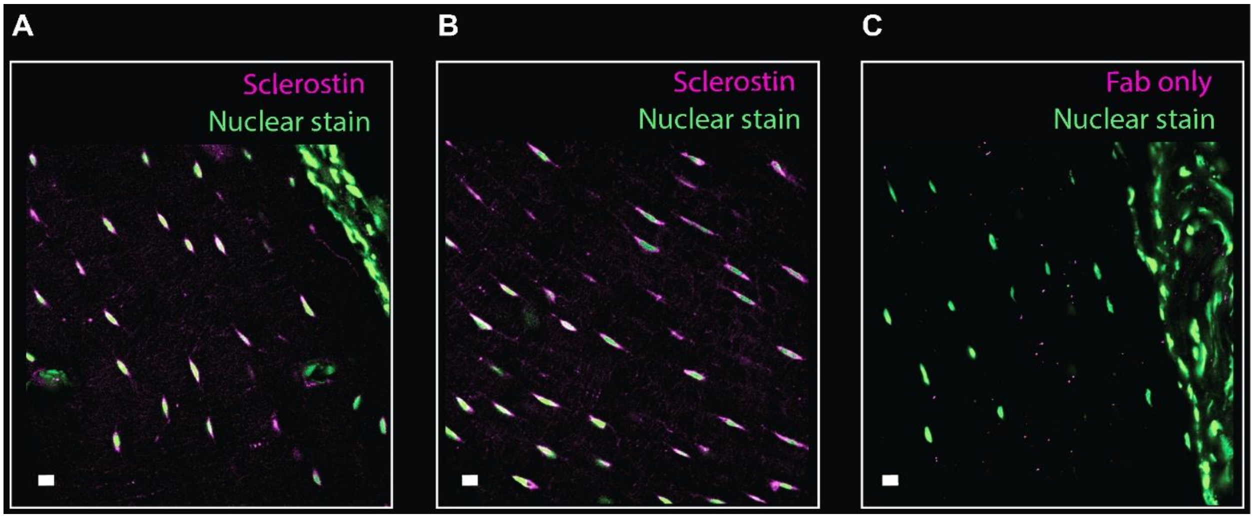

*Supplemental Figure 2: Protein labeling via nanobodies in mouse bone cryosections. A) Nanobodies anti-sclerostin staining. Signals can be observed in osteocytes. B) Nanobodies were amplified using Fab fragment anti-nanobodies conjugated with Alexa Fluor 647. C) Negative control using Fab fragment only without prior use of nanobodies. Images were acquired with a 40x objective using Zeiss LSM 800, located in CILS facility at Northeastern University Boston, MA. The scale represents 10 µm.*

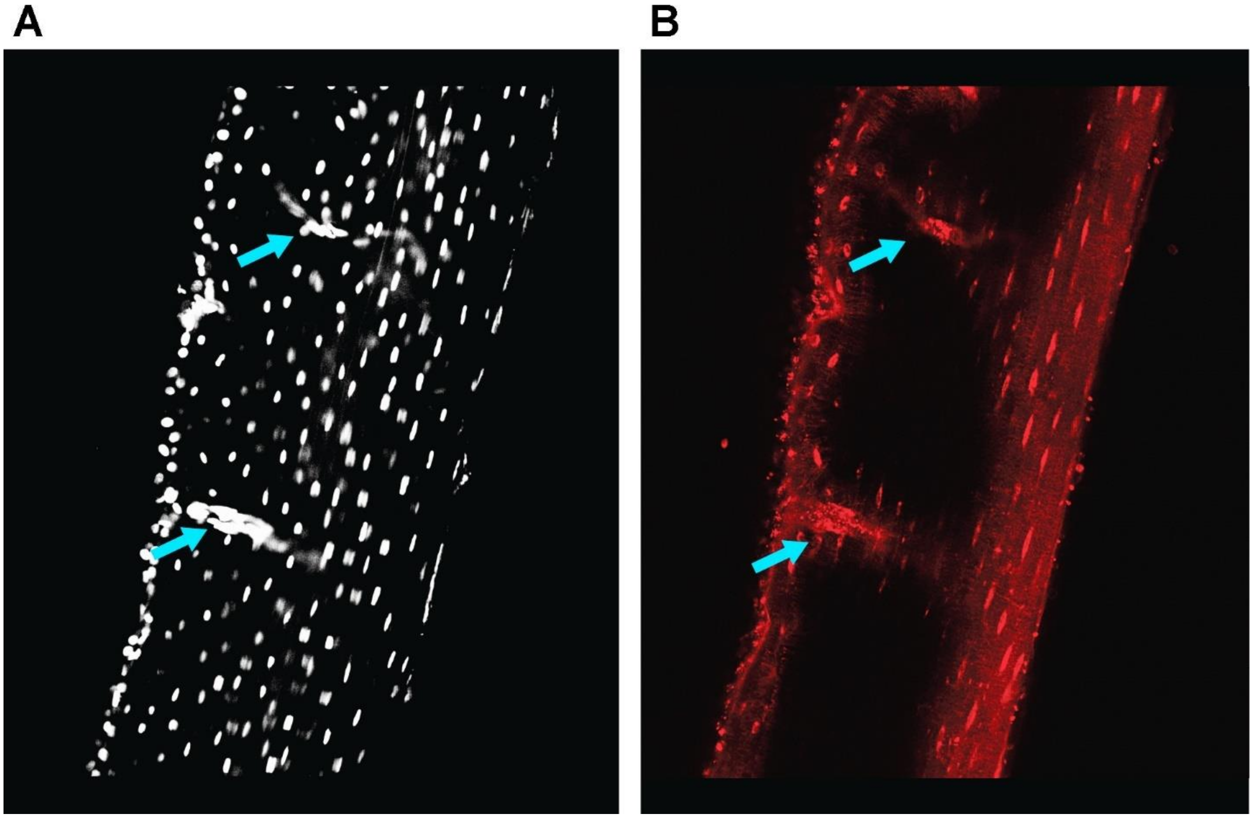

*Supplemental Figure 3: Blood vessels facilitate antibodies diffusion through the cortical bone. A) Longitudinal 2D view of a 3D mouse tibia midshaft showing signal from nuclear staining (Oxazole Yellow). Blue arrows point to two blood vessels in the cortical bone. B) Signal from anti-sclerostin antibodies shows penetration of the antibodies from both endosteal and periosteal surfaces, but also around the blood vessels. Lack of penetration if observed in the middle of the cortical thickness due to too short permeabilization step. Images were acquired using confocal microscopy.*

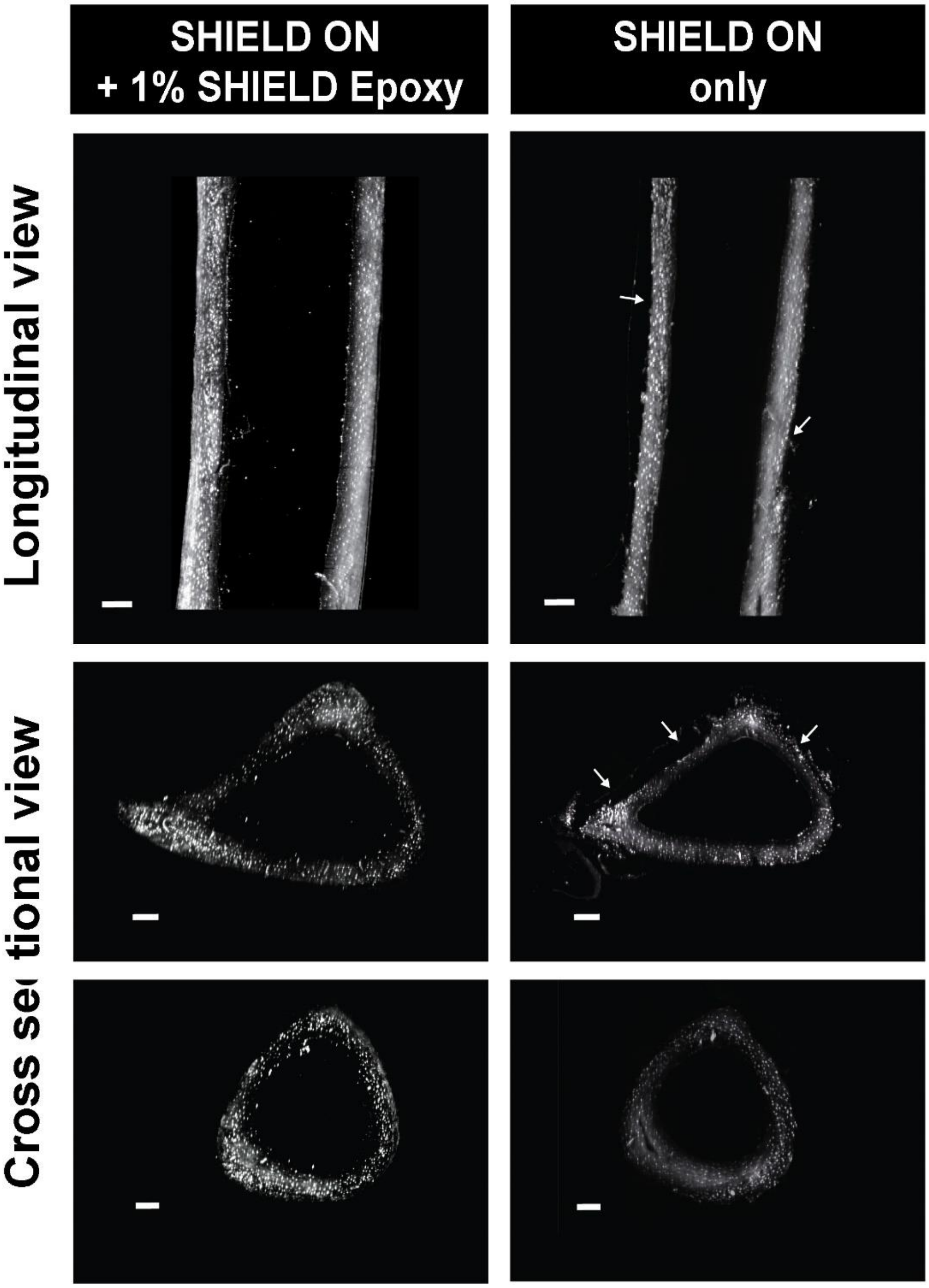

*Supplemental Figure 4 : Comparison of sample preservation and antibody (specific to sclerostin) penetration between two samples. The first sample was incubated in SHIELD ON Buffer with 1% of SHIELD Epoxy at 37°C overnight. The second sample was incubated in SHIELD ON Buffer at 37°C overnight. Both samples were then permeabilized with 10mg/ml of collagenase type II at 37°C for 6h. The addition of SHIELD Epoxy helped preserve samples edges from over-permeabilization. Over permeabilization regions can be observed for the sample that was incubated only in SHIELD ON buffer (white arrows).*

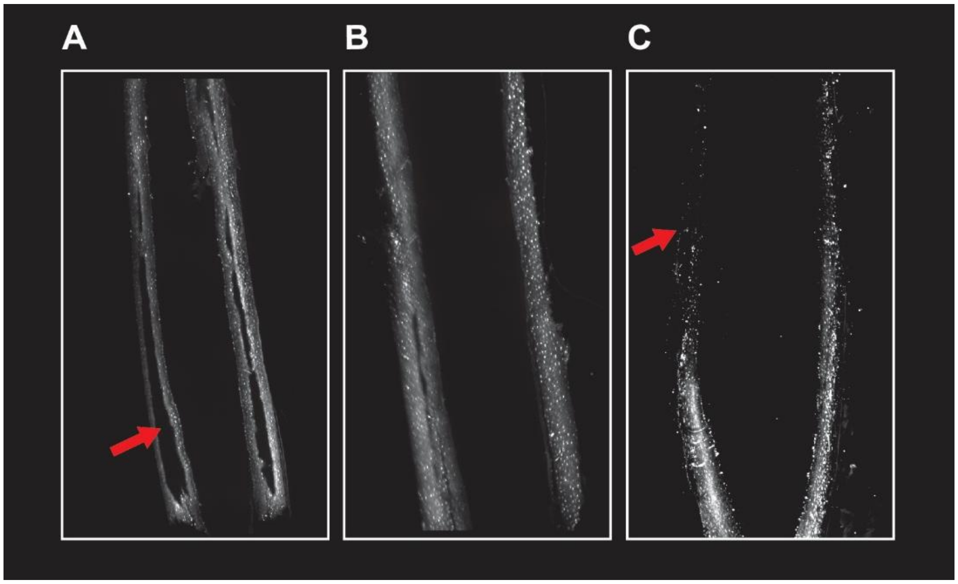

***Supplemental Figure 5: Optimization of the collagenase permeabilization step is critical to enable antibodies penetration without damaging the samples. A) Longitudinal 2D view of a 3D mouse tibia midshaft showing signal from anti-sclerostin antibodies. The red arrow points to a region of the sample where antibodies did not diffuse. B) Longitudinal 2D view of a 3D mouse tibia midshaft showing good penetration of the antibodies. C) Longitudinal 2D view of a 3D mouse tibia midshaft showing a damaged sample due to over permeabilization. The red arrow points to a region of a sample that was over permeabilized. The structure of the bone is disrupted. Images were acquired using lightsheet microscopy at 4x.***

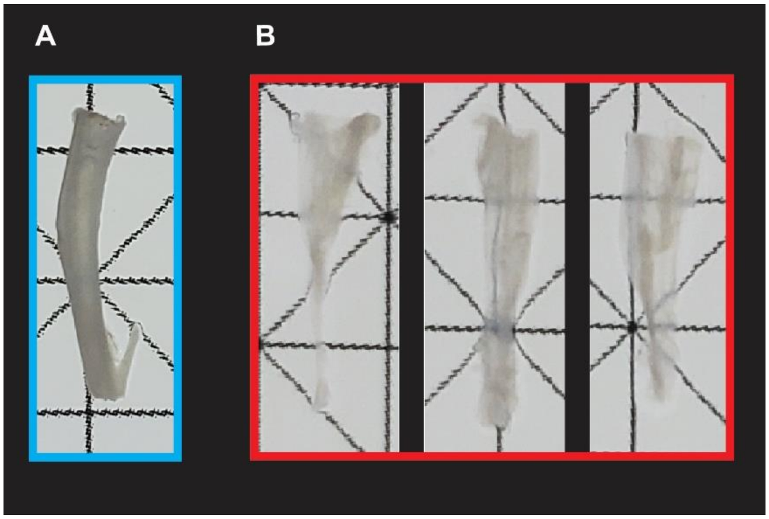

***Supplemental Figure 7: Example of over permeabilized mouse. A) Permeabilization step maintained the structure of the bone sample. This sample was SHIELD-preserved with addition of 1% SHIELD epoxy and permeabilized for 6h in 5ml of Collagenase type 2 (10 mg/ml) at 37°C. B) Examples of over-permeabilized SHIELD-preserved samples. Over permeabilization can lead to large damage of the bone. Samples will lose their structural integrity and will be difficult to manipulate. In this figure samples are shown in 1xPBS following the permeabilization step.***

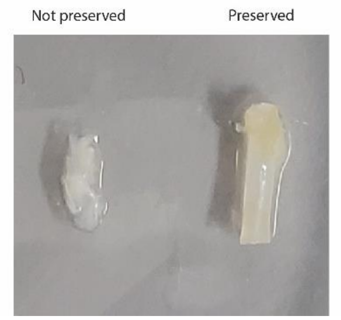

***Supplemental Figure 6: SHIELD-preserved versus non-preserved mouse femur midshaft after 3h of collagenase permeabilization. SHIELD allows preservation of the bone structure.***

### Supplemental methods

- **Sample mounting protocol**

The objective is to immobilize the sample for imaging within a media that has the same refractive index as the sample. Sample were mounted according to LifeCanvas Technologies' protocol available online.

*Gel preparation*: Mounting gel was prepared using 2% ultra-low melting point agarose (Sigma A5030) mixed with EasyIndex (LifeCanvas Technlogies, USA). For example, 0.6 g of agarose were added to 30 ml of EasyIndex. The tube was vortexed to ensure even distribution of the agarose. Gel was left at room temperature for an hour to allow agarose particles to be fully hydrated. The mounting gel was vortexed again to homogenize the solution and then aliquoted into 5ml tubes. A 5 ml tube of mounting gel was placed in a water bath at 90°C for at least an hour. During this time the tube was gently inverted several times to homogenize the gel. The tube was manipulated gently to avoid creating bubbles. After an hour, the gel was fully cleared and very few bubbles should be observed. If there were still bubbles, the gel would be placed back to the water bath for few minutes.

*Sample preparation:* Next, the temperature of the water bath was reduced to 80°C. In the meantime, an index-matched-bone sample was placed in a petri-dish with some EasyIndex. Using a light pad and a small lamp, the sample was inspected for any bubbles. Bubbles were removed from the medullary canal and from the surface of the bone using a 200 μl pipette.

*Sample holder preparation:* The sample holder was prepared by adding thermal tape or slices of sticky well plates covering each side. The gel was poured in the sample holder almost to the top. The 5ml tube containing the rest of the mounting gel was placed back in the water bath. The gel was inspected under light for any bubbles. Bubbles were pipetted out using a 200 μl pipette. Careful removal of any bubbles is critical to guarantee good image quality. After the gel was poured into the holder, manipulations were performed within 5min to prevent the gel from setting.

*Sample mounting:* Using a small spatula, the sample was slowly placed in the sample holder filled with gel. The bone sample was mounted longitudinally in the holder. The sample was inspected again under light to remove any bubbles created during the placement of the sample. The sample holder was placed at 4°C for 30 min on a flat surface. Next, a small top layer of mounting gel was added to the holder to fully cover the sample. The sample was returned to 4°C for 30min. Once set, the thermal tape was removed from the holder sides and the mounted sample was placed back in EasyIndex. The mounted sample was incubated overnight in Easyindex. Before imaging, mounted samples were rinsed in EasyIndex Matched Immersion oil for 10min.

- **Antibodies used for whole-mount immunolabeling:**

| Target | Antibodies |
|---|---|
| **Primaries** | |
| Connexin 43 | Rabbit IgG (3512S), Cell Signaling Technology |
| MMP9 | Goat IgG (AF909), RnD systems |
| Osteocalcin | Rabbit IgG (PA5-78870), Invitrogen |
| Osteopontin | Goat IgG (AF808), RnD systems |
| Sclerostin | Goat IgG (AF1589), RnD systems |
| Isotype control | Goat IgG Isotype Control (02-6202), Invitrogen |
| **Secondaries** | |
| Anti-Goat IgG | Donkey IgG Alexa Fluor Plus 555 (A32816), Invitrogen |
| Anti-Rabbit IgG | Donkey IgG SeTau-647 (AB_3095048), LifeCanvas Technologies |

***Table 1: List of Antibodies antibodies used for whole-mount immunolabeling.***

- **HCR-FISH probe sequences:**

| Pool name | Sequence |
|---|---|
| Mm_Sost_B1 | gAggAgggCAgCAAACggAAGCACAGAATCAGAATTGGCAGCTTT |
| Mm_Sost_B1 | ATCCACTATTACATCGGGCACTACGTAgAAgAgTCTTCCTTTACg |
| Mm_Sost_B1 | gAggAgggCAgCAAACggAATAGGTTGCCCCAGGATTCAACGACT |
| Mm_Sost_B1 | AAGTTGCTTTTCCCCACATCTGTAATAgAAgAgTCTTCCTTTACg |
| Mm_Sost_B1 | gAggAgggCAgCAAACggAACTTTGTGATCTGTGGGCAGGGTACT |
| Mm_Sost_B1 | CGCGGCAGCTGTACTCGGACACATTTAgAAgAgTCTTCCTTTACg |
| Mm_Sost_B1 | gAggAgggCAgCAAACggAACTGTCAGGAAGCGGGTGTAGTGCAG |
| Mm_Sost_B1 | CCGGCTTGGCGCTGCGGCATGGGCCTAgAAgAgTCTTCCTTTACg |
| Mm_Sost_B1 | gAggAgggCAgCAAACggAATGGCGTTGGGCAGCAGCCGCGCGGG |
| Mm_Sost_B1 | TCGGGCGCCACCACTTCACGCGCCCTAgAAgAgTCTTCCTTTACg |
| Mm_Sost_B1 | gAggAgggCAgCAAACggAACCGGGATGCAGCGGAAATCCGGTCC |
| Mm_Sost_B1 | GCTGCACCCGCTGCGCGCGGTAGCGTAgAAgAgTCTTCCTTTACg |
| Mm_Sost_B1 | gAggAgggCAgCAAACggAAACGCACCTTGCGCGAGCGCGGCGCC |
| Mm_Sost_B1 | GCGCTTGCACTTGCACGAGGCCACCTAgAAgAgTCTTCCTTTACg |
| Mm_Sost_B1 | gAggAgggCAgCAAACggAACTCCGACTGGTTGTGGAAGCGGGTG |
| Mm_Sost_B1 | CGCGGTCTCCGGCCCGAAGTCCTTGTAgAAgAgTCTTCCTTTACg |
| Mm_Sost_B1 | gAggAgggCAgCAAACggAACCGCGGCTTGCGACCCTTCTGCGGC |
| Mm_Sost_B1 | GTTGGCTTTGGCTCCCCGGGCGCCGTAgAAgAgTCTTCCTTTACg |
| Mm_Sost_B1 | gAggAgggCAgCAAACggAATGAAAACGAATCGGATCGCGCGGGG |
| Mm_Sost_B1 | CCCTGGCCTGGGCTGCAGGCTTTACTAgAAgAgTCTTCCTTTACg |
| Mm_Sost_B1 | gAggAgggCAgCAAACggAACTCCACACGGTCTGGAAAGTTTGGC |
| Mm_Sost_B1 | GACCTGCGGTCTCTACTGGGCTGGGTAgAAgAgTCTTCCTTTACg |
| Mm_Sost_B1 | gAggAgggCAgCAAACggAATCCTGGCCCGCGGGAACCCCACCCC |
| Mm_Sost_B1 | GGCAGAGTCTGGGACTCAAGCTTCCTAgAAgAgTCTTCCTTTACg |
| Mm_Sost_B1 | gAggAgggCAgCAAACggAAGGTCCGGCGAGGGTAGAAAGACCCC |
| Mm_Sost_B1 | GGTGGAAACACTGCCTTGTCCTGTATAgAAgAgTCTTCCTTTACg |
| Mm_Sost_B1 | gAggAgggCAgCAAACggAATTTCGTTCCACACTCCCTTCCCTTT |
| Mm_Sost_B1 | CTGTACGTCCATAACCAGTCCCAGGTAgAAgAgTCTTCCTTTACg |
| Mm_Sost_B1 | gAggAgggCAgCAAACggAAACATTTGGGTGGAAGGAGTAGATCT |
| Mm_Sost_B1 | AAACCCTATCTAGCCCACGCAGGCTTAgAAgAgTCTTCCTTTACg |
| Mm_Sost_B1 | gAggAgggCAgCAAACggAACACACTCAGTGGCCAGGTCAGGGTC |
| Mm_Sost_B1 | CCAAAAGAGAACCACGTAGCCCAACTAgAAgAgTCTTCCTTTACg |
| Mm_Sost_B1 | gAggAgggCAgCAAACggAAGGTCCCTATTTTACAAAGAAGACCG |
| Mm_Sost_B1 | CAATCCTTGGAATCTCAGCAGAGTTTAgAAgAgTCTTCCTTTACg |
| Mm_Sost_B1 | gAggAgggCAgCAAACggAATCTCTCACCAGTCTACACGGGGTAC |
| Mm_Sost_B1 | TCCCCTAACCCCTCCCCTGTTCTCCTAgAAgAgTCTTCCTTTACg |

| | |
|---|---|
| Mm_Sost_B1 | gAggAgggCAgCAAACggAATTCTAGGCGGTTGCCCACCACAATC |
| Mm_Sost_B1 | CGAGGCTGGGAGCCAACAAACAGCTTAgAAgAgTCTTCCTTTACg |
| Mm_Sost_B1 | gAggAgggCAgCAAACggAAGCAGATTTGAGAGGAAGGAGTGGGG |
| Mm_Sost_B1 | TTCCCTATCCCAGATATGGATTTGATAgAAgAgTCTTCCTTTACg |
| Mm_Sost_B1 | gAggAgggCAgCAAACggAATTACAATATGACAAGGCAGTTGGGG |
| Mm_Sost_B1 | CCTTAAACTGTTGTGTAGAAAATCCTAgAAgAgTCTTCCTTTACg |
| Mm_Sost_B1 | gAggAgggCAgCAAACggAAACTGGCAAGCCCAGTTTCCTCCAAC |
| Mm_Sost_B1 | CCTGGCAAGGGACAAGGATGGGAGGTAgAAgAgTCTTCCTTTACg |
| Mm_Sost_B1 | gAggAgggCAgCAAACggAACCGTGGGTGGCAGGCAGGAGGTGGT |
| Mm_Sost_B1 | ACGCTCTGTTTCTAGACAGAAATGTTAgAAgAgTCTTCCTTTACg |
| Mm_Sost_B1 | gAggAgggCAgCAAACggAAGTATAACACTTGCGCCCTCCGCCCC |
| Mm_Sost_B1 | GTGGCGCCTGACAGCTTCTCAGCATTAgAAgAgTCTTCCTTTACg |
| Mm_Sost_B1 | gAggAgggCAgCAAACggAAAGGTGTCTGGAAATGATTTACAAAA |
| Mm_Sost_B1 | ACAATTAAAATCTACACAGAAAGTATAgAAgAgTCTTCCTTTACg |
| Mm_Sost_B1 | gAggAgggCAgCAAACggAACGCCAAACTCACCAGCCCCAACCCC |
| Mm_Sost_B1 | TTGTGGATGAGTCTCACATGGAAAGTAgAAgAgTCTTCCTTTACg |
| Mm_Sost_B1 | gAggAgggCAgCAAACggAATAAATATGTCACTGAACTCTTAAAA |
| Mm_Sost_B1 | TTGGCATAAATAACTTAAATGAGAATAgAAgAgTCTTCCTTTACg |
| Mm_Sost_B1 | gAggAgggCAgCAAACggAAAACACTGCCTTTCTCTACAAGAAAA |
| Mm_Sost_B1 | CACACTTGTGCTTCACAAAGCGATATAgAAgAgTCTTCCTTTACg |
| Mm_Dmp1_B2 | CCTCgTAAATCCTCATCAAATCCATCGGGCTCTATTACTACTCCC |
| Mm_Dmp1_B2 | GCTAGCTACGCAAGCCTGCCATGGCAAATCATCCAgTAAACCgCC |
| Mm_Dmp1_B2 | CCTCgTAAATCCTCATCAAACACGAGACCAACACAAACTGTCGCA |
| Mm_Dmp1_B2 | TCTTGGCTGTGCCTCTCTCAGCAAAAAATCATCCAgTAAACCgCC |
| Mm_Dmp1_B2 | CCTCgTAAATCCTCATCAAAAGGACCTGCAAGGATGCCAAGTGAA |
| Mm_Dmp1_B2 | ATCGGCATGAGTCAGCAGGAGGGACAAATCATCCAgTAAACCgCC |
| Mm_Dmp1_B2 | CCTCgTAAATCCTCATCAAATAGCAGAGAAAGCCAGGTCTCCCCT |
| Mm_Dmp1_B2 | ACACAAATCGGCAGTAGTGGCAAGAAAATCATCCAgTAAACCgCC |
| Mm_Dmp1_B2 | CCTCgTAAATCCTCATCAAAGCCAGTTCCGTCGTATCGGGATAGC |
| Mm_Dmp1_B2 | GCAAACCCTTCAAGGATATACCAGCAAATCATCCAgTAAACCgCC |
| Mm_Dmp1_B2 | CCTCgTAAATCCTCATCAAACAGCAGCGGCAGCAGCTCAGTCCAT |
| Mm_Dmp1_B2 | GAAATCTGCAGTTCAGTTCACAGGAAAATCATCCAgTAAACCgCC |
| Mm_Dmp1_B2 | CCTCgTAAATCCTCATCAAAATGCGATTCCTCTACCTGCGTTGTC |
| Mm_Dmp1_B2 | AAGGAGAATGACAGTCTTCATATTGAAATCATCCAgTAAACCgCC |
| Mm_Dmp1_B2 | CCTCgTAAATCCTCATCAAAAACTGGGAGAGCACAGGACAGCCCC |
| Mm_Dmp1_B2 | TTCAGATTCAGTATTGTGGTATCTGAAATCATCCAgTAAACCgCC |
| Mm_Dmp1_B2 | CCTCgTAAATCCTCATCAAACAAATCACCCGTCCTCTCTTCAGAG |
| Mm_Dmp1_B2 | GTTCGTGGGTGGTGGTGGTGACCCAAAATCATCCAgTAAACCgCC |

| | |
|---|---|
| Mm_Dmp1_B2 | CCTCgTAAATCCTCATCAAAAGCTTGACTTTCTTCTGATGACTCA |
| Mm_Dmp1_B2 | GTCACTATTTGCCTGTCCCTCTGGGAAATCATCCAgTAAACCgCC |
| Mm_Dmp1_B2 | CCTCgTAAATCCTCATCAAACTCTCCAGATTCACTGCTGTCCGTG |
| Mm_Dmp1_B2 | GTACTGGCCTCTGTCGTAGCCCAGCAAATCATCCAgTAAACCgCC |
| Mm_Dmp1_B2 | CCTCgTAAATCCTCATCAAAACTCTTAGAGAGTCCACCAGCCGGT |
| Mm_Dmp1_B2 | ATCCTCCTTATCGGCGCCGGTCCCCAAATCATCCAgTAAACCgCC |
| Mm_Dmp1_B2 | CCTCgTAAATCCTCATCAAAGGTATCATCTCCACTGTCGTCTTCA |
| Mm_Dmp1_B2 | CCCTAGATCATTGTCCTCATCGCCAAAATCATCCAgTAAACCgCC |
| Mm_Dmp1_B2 | CCTCgTAAATCCTCATCAAAGGGTCCTCCCCACTGTCCTTCTTCG |
| Mm_Dmp1_B2 | GGAGTCCTCATCACTGTCCAGTTTGAAATCATCCAgTAAACCgCC |
| Mm_Dmp1_B2 | CCTCgTAAATCCTCATCAAAGTCTTCACTGGACTGTGTGGTGTCT |
| Mm_Dmp1_B2 | TTGGGCACTGTTTTCTTGAGAGGTGAAATCATCCAgTAAACCgCC |
| Mm_Dmp1_B2 | CCTCgTAAATCCTCATCAAAGTCGTGGTCTTTGCTGTCGCTGGGG |
| Mm_Dmp1_B2 | AGGCCGGCTGTCTGCCTCATCCTCAAAATCATCCAgTAAACCgCC |
| Mm_Dmp1_B2 | CCTCgTAAATCCTCATCAAAACTGTCCTGAGTGGAGTCGCCTGCC |
| Mm_Dmp1_B2 | ACCTCCCACCCGCTGTTCCTCACTCAAATCATCCAgTAAACCgCC |
| Mm_Dmp1_B2 | CCTCgTAAATCCTCATCAAAAACCGTCCCCGTGGCTACTCTCCCC |
| Mm_Dmp1_B2 | TCTGCATCCCTTCATCATCGAACTCAAATCATCCAgTAAACCgCC |
| Mm_Dmp1_B2 | CCTCgTAAATCCTCATCAAACGCTCCTGGTACTCTCGGGGTCGTC |
| Mm_Dmp1_B2 | CGCTGCTCATTCTGGCGTGGCCTCGAAATCATCCAgTAAACCgCC |
| Mm_Dmp1_B2 | CCTCgTAAATCCTCATCAAACTTTAGATTCTTCCGACCTGATACC |
| Mm_Dmp1_B2 | CCTGAGTGCTCGTGGGCTCGTGGTCAAATCATCCAgTAAACCgCC |
| Mm_Dmp1_B2 | CCTCgTAAATCCTCATCAAAATTCCACAGACTGGCTGTCATCTGA |
| Mm_Dmp1_B2 | ACCTTCTGAAGGACTTCCTGCTTGAAAATCATCCAgTAAACCgCC |
| Mm_Dmp1_B2 | CCTCgTAAATCCTCATCAAACTCTGTAGTCTTCCTCAGAGACGTG |
| Mm_Dmp1_B2 | CCCTGCTGTTGCTGTCAGTAAGCTCAAATCATCCAgTAAACCgCC |
| Mm_Dmp1_B2 | CCTCgTAAATCCTCATCAAATATCCTCCGTGGAGTCGCTCTGGGT |
| Mm_Dmp1_B2 | CGCTCCTGCTTTCCTCCTTGGAGGCAAATCATCCAgTAAACCgCC |
| Mm_Dmp1_B2 | CCTCgTAAATCCTCATCAAAGGCTCTCGGCTGTGTCCTCCTGAGA |
| Mm_Dmp1_B2 | GCCCCTCTGGGCTATCTTCCTGGGAAAATCATCCAgTAAACCgCC |
| Mm_Dmp1_B2 | CCTCgTAAATCCTCATCAAACTTCGCTGGACTCACTGCTGGGGTC |
| Mm_Dmp1_B2 | TGCTTTCCTGGGATGGCTCACCAGCAAATCATCCAgTAAACCgCC |
| Mm_Dmp1_B2 | CCTCgTAAATCCTCATCAAATGGTCACCCCTTCCTGAGATTCGCT |
| Mm_Dmp1_B2 | TATCTGGGTTGTCACCCCTGGACTCAAATCATCCAgTAAACCgCC |
| Mm_Dmp1_B2 | CCTCgTAAATCCTCATCAAACTTCTTGGTCTCCTGCCTGACTTGT |
| Mm_Dmp1_B2 | GGCTGTCCTCCTCACTGGACTCACTAAATCATCCAgTAAACCgCC |
| Mm_Dmp1_B2 | CCTCgTAAATCCTCATCAAAGGCTTTCTGAGCTGGAGAATGTGTT |
| Mm_Dmp1_B2 | CGCTGTCAGCTTGCTCCTCGGTGGAAAATCATCCAgTAAACCgCC |
| Mm_Dmp1_B2 | CCTCgTAAATCCTCATCAAACGGAGAGGCTGAGGCTCTCGTTGGA |

| | |
|---|---|
| Mm_Dmp1_B2 | CATCCTGGGCCGACTCCTGACTCTCAAATCATCCAgTAAACCgCC |
| Mm_Dmp1_B2 | CCTCgTAAATCCTCATCAAAGCAGGCCTTCCTGGCTGGAGCTGTC |
| Mm_Dmp1_B2 | TGCTCTCAGTGGATGCGCTCTGGGAAAATCATCCAgTAAACCgCC |
| Mm_Dmp1_B2 | CCTCgTAAATCCTCATCAAACCTGCTCAGACTGGCTCTCCTGGCT |
| Mm_Dmp1_B2 | AGTCACTGTCTTCCTCAGAACGGCTAAATCATCCAgTAAACCgCC |
| Mm_Dmp1_B2 | CCTCgTAAATCCTCATCAAACTTCTTTGGATCGGCTACTGTCCTG |
| Mm_Dmp1_B2 | TGGAAGCGCTCCCTGTGGAGTTGCTAAATCATCCAgTAAACCgCC |
| Mm_Dmp1_B2 | CCTCgTAAATCCTCATCAAATCTTGGGACGGATGTCCTCCTCGCT |
| Mm_Dmp1_B2 | TTAGTTTCCTACTGTCAGCTTCCATAAATCATCCAgTAAACCgCC |
| Mm_Dmp1_B2 | CCTCgTAAATCCTCATCAAATGGGTTTGTTGTGGTAAGCATCAAC |
| Mm_Dmp1_B2 | GACAGTCATTGTCATCTTGGTCCCCAAATCATCCAgTAAACCgCC |
| Mm_Dmp1_B2 | CCTCgTAAATCCTCATCAAAAGACAAGCTAATGCTAGTAGCCGTC |
| Mm_Dmp1_B2 | GACTCCGTCCTGTGAGAGCCATTTCAAATCATCCAgTAAACCgCC |
| Mm_Dmp1_B2 | CCTCgTAAATCCTCATCAAAGGGGAAAACAAACAAACAAACAAAA |
| Mm_Dmp1_B2 | AGTGTCGTGTCCAACAGTTCCTCACAAATCATCCAgTAAACCgCC |
| Mm_Dmp1_B2 | CCTCgTAAATCCTCATCAAATAAGGAGTGGGAGACACCCTGGAAA |
| Mm_Dmp1_B2 | TGACATCATCCCACGTACTTAAGCCAAATCATCCAgTAAACCgCC |
| Mm_Dmp1_B2 | CCTCgTAAATCCTCATCAAATGTGTTCTTCTCTCACCATGTGTGC |
| Mm_Dmp1_B2 | CTGTGAAGCGGCTGTGCACGTGATCAAATCATCCAgTAAACCgCC |
| Mm_Dmp1_B2 | CCTCgTAAATCCTCATCAAACACTGTCAGGTTGGTGAACCAGAGC |
| Mm_Dmp1_B2 | GTGCCTGGTGTGTACCAGAACCTCAAAATCATCCAgTAAACCgCC |
| Mm_Dmp1_B2 | CCTCgTAAATCCTCATCAAAACTGAGCCTGAAGCACCACCACCCC |
| Mm_Dmp1_B2 | CCTTGGAACAAGCTCCTCCGGAGCAAAATCATCCAgTAAACCgCC |
| Mm_Dmp1_B2 | CCTCgTAAATCCTCATCAAACCTTTAATGTGTTACTTTTGAGAGT |
| Mm_Dmp1_B2 | CCAGCTCCAGGGGCTCCTTCTTTGTAAATCATCCAgTAAACCgCC |
| Mm_Dmp1_B2 | CCTCgTAAATCCTCATCAAAACCCTCCAGGAGCCAAAGCAATGTA |
| Mm_Dmp1_B2 | GGCAGGGGCTTGCTGCAGTAAGTTCAAATCATCCAgTAAACCgCC |
| Mm_Dmp1_B2 | CCTCgTAAATCCTCATCAAAATTCCCTGTGGTACTCGCCTCAGCT |
| Mm_Dmp1_B2 | AAACTGGTCACTTCCTGTCCTGCTCAAATCATCCAgTAAACCgCC |
| Mm_Dmp1_B2 | CCTCgTAAATCCTCATCAAACTGTGGAACCTCTGATTCTCCAAAG |
| Mm_Dmp1_B2 | GTAACCCTCCAGCTCCAGGCTTTGCAAATCATCCAgTAAACCgCC |
| Mm_Dmp1_B2 | CCTCgTAAATCCTCATCAAAATATTGTCTTCAATGGAGAAATCCT |
| Mm_Dmp1_B2 | CTGCTTATCTCAAGAATAAATAGAGAAATCATCCAgTAAACCgCC |
| Mm_Dmp1_B2 | CCTCgTAAATCCTCATCAAAAAATATTTAATTGCATACGATTTGA |
| Mm_Dmp1_B2 | TATCATTTGAGAATGCCTTTACGTAAAATCATCCAgTAAACCgCC |
| Mm_Dmp1_B2 | CCTCgTAAATCCTCATCAAATATGTTATCCAAGATATCCCATTGG |
| Mm_Dmp1_B2 | TCAATAAGGTAAATTCGTTGGGAAAAAATCATCCAgTAAACCgCC |
| Mm_Dmp1_B2 | CCTCgTAAATCCTCATCAAAATGTTTACAGAAGGCCCACAAAAGC |
| Mm_Dmp1_B2 | TTCTTCCTCCTCCTCATCATTGAGGAAATCATCCAgTAAACCgCC |

| | |
|---|---|
| Mm_Dmp1_B2 | CCTCgTAAATCCTCATCAAACACTGCACATAAATAATCTGGAAAA |
| Mm_Dmp1_B2 | ATAGAAAGGACTCAGGGATAGTATTAAATCATCCAgTAAACCgCC |
| Mm_Dmp1_B2 | CCTCgTAAATCCTCATCAAAACTTGAGCCCTATGTGCAGAATTTT |
| Mm_Dmp1_B2 | GAAAAGATAAAGAAACAACAACAAAAAATCATCCAgTAAACCgCC |

***Table 2: List of probes designed for whole-mount mRNA labeling.***
